# Consequences of alanine-126 mutations in helix-3 on structure and functions of Rad6 E2 ubiquitin-conjugating enzymes

**DOI:** 10.1101/2021.10.12.464121

**Authors:** Prakash K. Shukla, Dhiraj Sinha, Andrew M. Leng, Jesse E. Bissell, Shravya Thatipamula, Rajarshi Ganguly, Jack J. Skalicky, Dennis C. Shrieve, Mahesh B. Chandrasekharan

**Author notes:** Corresponding author Mahesh B. Chandrasekharan, 2000 Circle of Hope Room 3715, Huntsman Cancer Institute, University of Utah School of Medicine, Salt Lake City, UT 84112, USA.

## Abstract

Rad6, an E2 ubiquitin-conjugating enzyme conserved from yeast to humans, functions in transcription, genome maintenance and proteostasis. The contributions of many conserved secondary structures of Rad6 and its human homologs UBE2A and UBE2B to their biological functions are not understood. A mutant *RAD6* allele with a missense substitution at alanine-126 (A126) of helix**-**3 that causes defects in telomeric gene silencing, DNA repair and protein degradation was reported over two decades ago. Here, using a combination of genetics, biochemical, biophysical, and computational approaches, we discovered that helix**-**3 A126 mutations compromise the ability of Rad6 to ubiquitinate target proteins without disrupting interactions with partner E3 ubiquitin-ligases that are required for their various biological functions *in vivo*. Explaining the defective *in vitro* or *in vivo* ubiquitination activities, molecular dynamics simulations and NMR showed that helix**-**3 A126 mutations cause local disorder of the catalytic pocket of Rad6 in addition to disorganizing the global structure of the protein to decrease its stability *in vivo*. We also show that helix**-**3 A126 mutations deform the structures of UBE2A and UBE2B, the human Rad6 homologs, and compromise the *in vitro* ubiquitination activity and folding of UBE2B. In summary, our studies reveal that the conserved helix**-**3 is a crucial structural constituent that controls the organization of catalytic pockets, enzymatic activities, and biological functions of the Rad6-family E2 ubiquitin-conjugating enzymes.

## Introduction

Ubiquitination, the covalent post-translational modification of proteins by the highly conserved 76 amino-acid protein ubiquitin (Ub), controls many aspects of cellular function (1,2). Ubiquitination is a three-step process: First, an E1 ubiquitin-activating enzyme uses ATP to activate ubiquitin. Second, the E1 ubiquitin-activating enzyme attaches the ubiquitin onto the active-site cysteine of an E2 ubiquitin-conjugating enzyme (henceforth referred to as an E2 enzyme) (1,3). Third, an E3 ubiquitin-ligase (henceforth referred to as E3 ligase) and the ubiquitin-charged E2 enzyme target a substrate protein to catalyze the formation of an isopeptide bond between the C-terminus of ubiquitin and a nucleophile, which is typically a lysine side chain on the substrate protein (4). Monoubiquitination or addition of just one ubiquitin moiety is important during transcription and DNA repair (5-7). Ubiquitin contains seven lysine residues, which are targeted for cycles of ubiquitin addition to form polyubiquitin chains. Polyubiquitination through ubiquitin lysine-48 (K48) generally targets proteins for proteasomal degradation, whereas K63-linked ubiquitin chains regulate signal transduction and endocytosis (1,8). Misregulation of ubiquitination is associated with numerous diseases ranging from neurological disorders to cancers (9-14).

E2 enzymes are central players in protein ubiquitination (15,16). Humans express ∼35 and *Saccharomyces cerevisiae* or budding yeast express 12 E2 enzymes. The E2 enzymes contain a distinctive core catalytic domain of about 150 amino acids called the UBC fold (17), which is comprised of four α-helices, four β-sheets (also called a β-meander), and a conserved active-site cysteine (18). Additional residues also have roles in the catalytic functions of E2 enzymes: one is a conserved asparagine in a ‘flap’ or loop region present close to the active-site cysteine, which is part of the HxN triad that is proposed to aid in localizing the target lysine (19) or the active site (20) and in stabilizing the oxyanion formed in the reaction intermediate during the nucleophilic attack (21). Another is the “gateway residue”, which is a conserved serine or aspartate present in a loop that forms the opening of the E2 active-site cleft and implicated in regulation of E2 activity (15,22). Some E2 enzymes also contain internal and/or N- or C-terminal extensions to the UBC fold that have regulatory functions (14).

Functions for various secondary structures within the UBC domain of E2 enzymes have also been reported (15,23): the N-terminal helix-1 in some E2s is an E1 or E3 binding surface. Loops in the front face close to the catalytic pocket of E2s are functionally important in binding the RING domain of E3 ligases. The vast majority of the non-covalent interactions of E2 enzymes with ubiquitin, partner E3 ubiquitin-ligases, or other regulatory factors involve a so-called “backside” surface that is located on the face opposite from the catalytic pocket and made up of residues of the four β-sheets, the C-terminal end of helix-1, the intervening loops and the C-terminal end of helix-4 (23-27). Despite these many structure-function studies, the roles for other regions within E2 enzymes, such as helix-3, remain not known.

Rad6 (<u>Rad</u>iation sensitive <u>6</u>) is an E2 enzyme in budding yeast that has well-established functions in transcription, DNA repair, and protein homeostasis that are accomplished through its interactions with different partner E3 ligases (Figure 1a): Rad6 interacts with the Rad18 E3 ligase to monoubiquitinate PCNA and activate translesion DNA repair following DNA damage (28,29). During transcription and other nuclear processes, Rad6 interacts with the Bre1 E3 ligase and the adapter protein Lge1 to monoubiquitinate histone H2B at K123 (H2BK123) (30-33), which in turn participates in the *trans*-histone regulation of methylation of histone H3 at K4 and K79 (34-37). Rad6 partners with the E3 ligase Ubr2 and the adapter protein Mub1 to polyubiquitinate phosphorylated Sml1, a ribonucleotide reductase inhibitor, resulting in its proteasomal degradation upon recovery from DNA damage (38). The Rad6-Ubr2 E2-E3 complex is also reported to regulate Rpn4 and Dsn1 protein levels via ubiquitination (39,40). In the N-end rule pathway of targeted proteolysis, Rad6 and the Ubr1 E3 ligase polyubiquitinate various proteins to target them for degradation by the proteasome machinery (41-43).

**Fig. 1.**
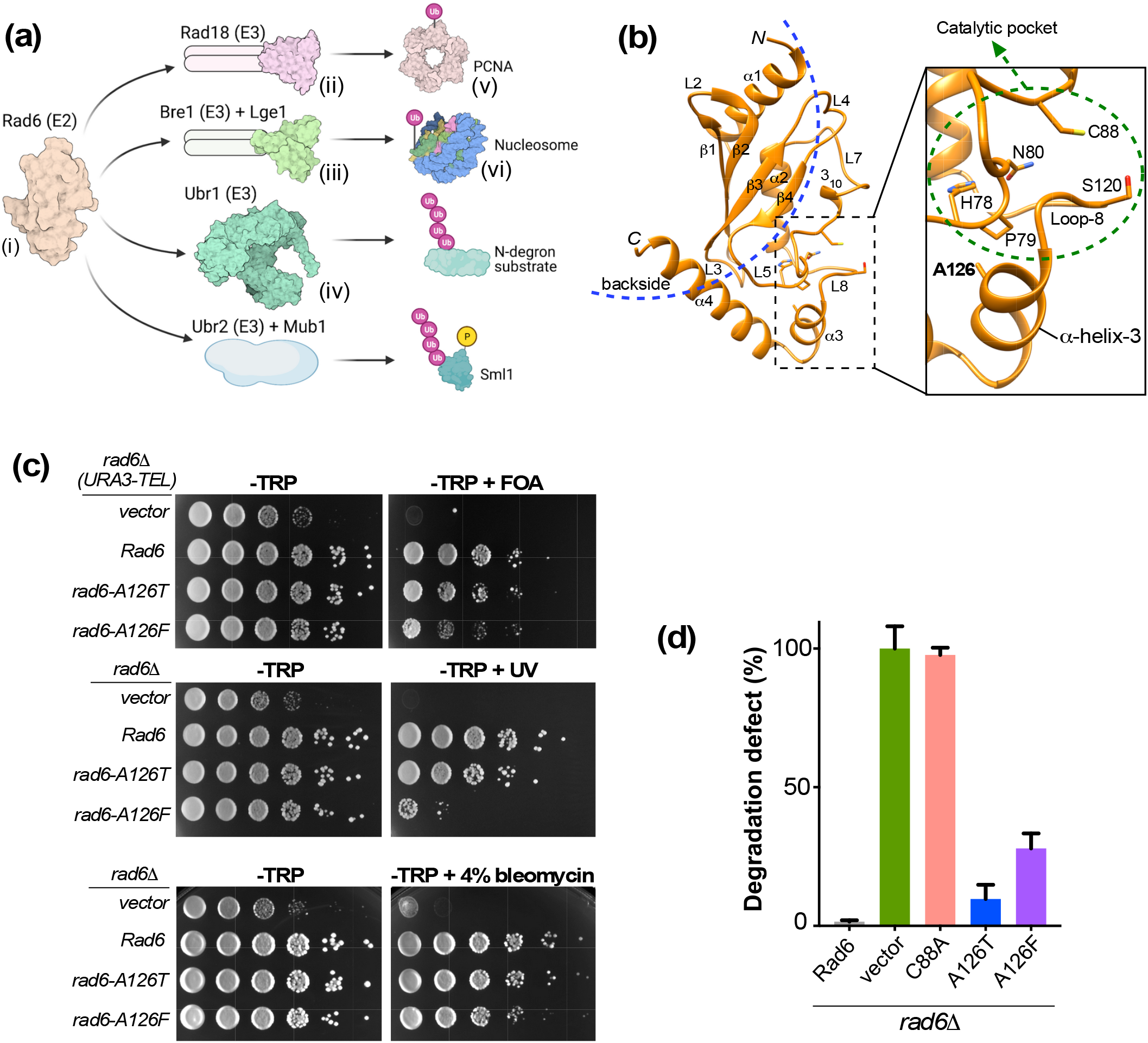
Rad6 A126 mutations cause defects in telomeric gene silencing, DNA repair and protein degradation in yeast. (a) Illustration shows Rad6 and its partner E3 ubiquitin-ligases (Rad18, Bre1, Ubr1 or Ubr2) involved in mono- or poly-ubiquitination of the indicated target proteins. Crystal structure data were used in depictions of *i)* Rad6 (PDB ID: 1AYZ), *ii*, the Rad18 RING domains (PDB ID: 2Y43), *iii*, the Bre1 RING domains (PDB ID: 4R7E), *iv*, Ubr1 (PDB ID: 7MEX), *v*, PCNA (PDB ID: 6D0R), and *vi*, yeast nucleosome (PDB ID: 1ID3). Abbreviations: Ub, ubiquitin; P, phosphate. (b) Ribbon representation of Rad6. Secondary structures including α-helices, 3_10_-helix, β- sheets, and intervening loops (L) are labeled. The region to the left of the *blue dotted arc* is the backside region of Rad6 comprised of residues in the β-sheets, intervening loops, and in the C-terminal ends of helices 1 and 4. Zoomed image shows the location of A126 in helix-3 and its spatial proximity to the catalytic pocket (encircled by *green dotted line*), which is comprised of the HPN motif, active-site C88 and the gateway residue S120 in loop-8. (c) *Top panel*, Growth assay for telomeric gene silencing was conducted by spotting ten-fold serial dilutions of indicated strains on synthetic medium lacking tryptophan (-TRP) or containing 5-fluoroorotic acid (-TRP+FOA). *Middle panel*, Growth assay conducted by spotting a ten-fold serial dilution of the indicated strains on medium containing tryptophan with or without exposure to UV light. *Bottom panel*, Growth assay was conducted by spotting a ten-fold serial dilution of the indicated strains on medium containing tryptophan with or without 4% bleomycin. (d) β- galactosidase activity was measured in extracts of *rad6Δ* strains co-transformed with the plasmid for expression of N-end rule degradation reporter (R-β-gal) and empty vector or constructs to express wild-type Rad6 or indicated mutants. Absence of Rad6 (*rad6Δ*) or its activity (*rad6-C88A)* stabilizes the reporter yielding high β-gal activity or 100% degradation defect. In contrast, complete degradation of the reporter occurs in the presence of wild-type Rad6 or zero degradation defect. Plotted are means ± standard error from three replicate assays.

Structure-function studies have revealed that the N-terminal nine residues of helix-1 and amino acids 150-153 of Rad6 are necessary for its interaction with the Ubr1 E3 ligase (44). Residues 141-149 at the C-terminus and residues 10-22 at the N-terminus of Rad6 are necessary for binding to the Rad18 E3 ligase (45). A non-RING domain N-terminal region (amino acids 1-210) of the Bre1 E3 ligase binds the backside of Rad6 (46,47). Rad6 possess a 23 amino-acid acidic tail that is important for its enzymatic activity *in vitro* and *in vivo* (48-50). Phosphorylation of gateway residue serine-120 (S120) of Rad6 occurs *in vivo* and regulates monoubiquitination of H2BK123 (51). Based on the studies on Rad6 and other E2 enzymes (18,19,21,51,52), the active-site C88, S120 in the active-site cleft or gate, and H78 and N80 in the HPN motif together constitute the catalytic pocket of Rad6 (Figure 1b). Low-affinity non-covalent interactions of ubiquitin with the backside of Rad6 was reported to influence its processivity (24). Collectively, these studies have defined the role(s) of various residues and secondary structures within Rad6 to its protein-protein interactions and enzymatic functions; however, the contributions of other regions of the UBC domain to overall structure and enzymatic functions are not fully understood.

Nearly two decades ago, Liebman and colleagues reported the isolation and initial characterization of a missense threonine substitution at alanine (A) 126 in helix-3 of yeast Rad6. This mutant was defective in telomeric gene silencing and other functions of Rad6 (53). Here, using a multidisciplinary approach, we examined the contributions of A126 in helix-3 to the biological functions of Rad6. Mutations at A126 adversely affected the Rad6-mediated mono- and polyubiquitination of substrate proteins that regulate telomeric gene silencing, DNA repair and protein degradation. Using molecular dynamics (MD) simulations and NMR, we show that mutations in A126 cause disorganization of local structure of the catalytic pockets as well as global structure of Rad6 that inhibit enzymatic activity and/or compromise protein stability in an E3 ligase-independent fashion. Our investigations further show that A126 mutation(s) in helix-3 also disrupt the structure and enzymatic activity of human Rad6 homologs. Overall, our studies reveal that the conserved helix-3 is a crucial structural constituent of yeast and human Rad6 E2 ubiquitin-conjugating enzymes.

## Results

### A126 mutations disrupt telomeric gene silencing, DNA repair and protein degradation functions of Rad6

A screen for *RAD6* alleles that cause defects in telomeric gene silencing identified a mutant with a threonine substitution at A126 of Rad6 (53). The *rad6-A126T* allele also conferred sensitivity to ultraviolet (UV) irradiation, indicating impaired DNA repair function, and was also shown to be defective in N-end rule protein degradation (53). A126 is in helix-3 of Rad6, close to the catalytic pocket (Figure 1b). To investigate how mutations in A126 residue influence the biological functions of Rad6 in telomeric gene silencing, DNA repair and protein degradation, we created yeast constructs for expression of the rad6-A126T mutant, and of a Rad6 protein with a bulkier phenylalanine (F) substitution (rad6-A126F).

To test the effect of these mutations on telomeric gene silencing, we transformed the constructs into a *rad6* null mutant yeast strain (*rad6Δ*) that harbors a silencing reporter gene (*URA3*) integrated near the left end of chromosome VII. As controls, either the empty vector or a plasmid for expression of wild-type Rad6 were transformed into this reporter strain. Transcriptional repression of the telomere-proximal *URA3* reporter occurred in the presence of wild-type Rad6, which allowed yeast cells to survive on a media containing the counterselection agent 5-fluoroorotic acid (5FOA)(54), whereas the absence of Rad6 resulted in the transcriptional activation and production of the URA3 enzyme, which converts 5FOA to toxic 5-fluorouracil, inhibiting growth (Figure 1c, top panels). Consistent with the previous report (53), expression of the rad6-A126T mutant resulted in slower growth on 5FOA medium compared to the control strain that expressed wild-type Rad6 (Figure 1c), indicating activation of the *URA3* reporter and a telomeric silencing defect. The rad6-A126F mutant showed a more drastic growth retardation on 5FOA medium than the rad6-A126T mutant strain (Figure 1c), revealing that this mutation causes a severe telomeric gene silencing defect. Next, we examined the growth of Rad6 A126 mutant strains following exposure to DNA damaging agents: UV or the radiomimetic drug bleomycin (55). The rad6-A126T mutant showed a subtle slow growth defect and the rad6-A126F mutant had a more severe growth retardation upon UV irradiation or bleomycin treatment when compared to the control strain that expresses wild-type Rad6 (Figure 1c, middle and lower panels). These results suggest that mutations in A126 residue impair the DNA repair functions of Rad6.

Proteins with N-terminal arginine are degraded by the N-end-rule pathway, where polyubiquitination by Rad6 precedes proteasome-mediated proteolysis (56). To measure the N-end rule activity, extracts were prepared for enzyme assays from strains expressing either wild-type or mutant Rad6 along with the reporter beta-galactosidase protein with arginine as the N-terminal amino acid (R-β-Gal) (57). Low or no activity was obtained for R-β-Gal in extracts prepared from the strain with wild-type Rad6 (Figure 1d). In contrast, extracts prepared from strains lacking Rad6 (*rad6Δ*) or expressing the catalytic-dead mutant (rad6-C88A) yielded very high activity compared to the control strain with wild-type Rad6 (Figure 1d), indicating stabilization of R-β-Gal levels and a defect in the N-end-rule degradation. Extracts prepared from rad6-A126T and rad6-A126F mutant strains showed higher galactosidase activity than the control strain with wild-type Rad6 (Figure 1d), indicating that these mutants are also defective in the N-end rule protein degradation process. Collectively, our results confirmed that mutations at A126 compromise the functions of Rad6 in telomeric gene silencing, DNA repair and targeted proteolysis.

### A126 mutations adversely affect Rad6-mediated mono- or poly-ubiquitination of substrate proteins *in vivo*

Next, we asked whether Rad6 A126 mutations disrupt ubiquitination of substrate proteins histone H2B and PCNA or Sml1, which are involved in telomeric gene silencing and DNA repair, respectively (28,34,38). A complex of Rad6, Bre1 and Lge1 catalyzes histone H2BK123 monoubiquitination (H2Bub1) (30-32), which regulates methylation (me) of histone H3K4 (34,36). Decreased trimethylation of H3K4 (H3K4me3) causes telomeric gene silencing defect (37,58,59). Therefore, we measured the steady-state levels of H2Bub1 and H3K4me in *rad6-A126T* and *rad6-A126F* strains using immunoblotting. When expressed in the *rad6Δ* strain, both rad6-A126T and rad6-A126F mutants caused an apparent complete loss of H2Bub1 (Figure 2a, lanes 3-4).

**Fig. 2.**
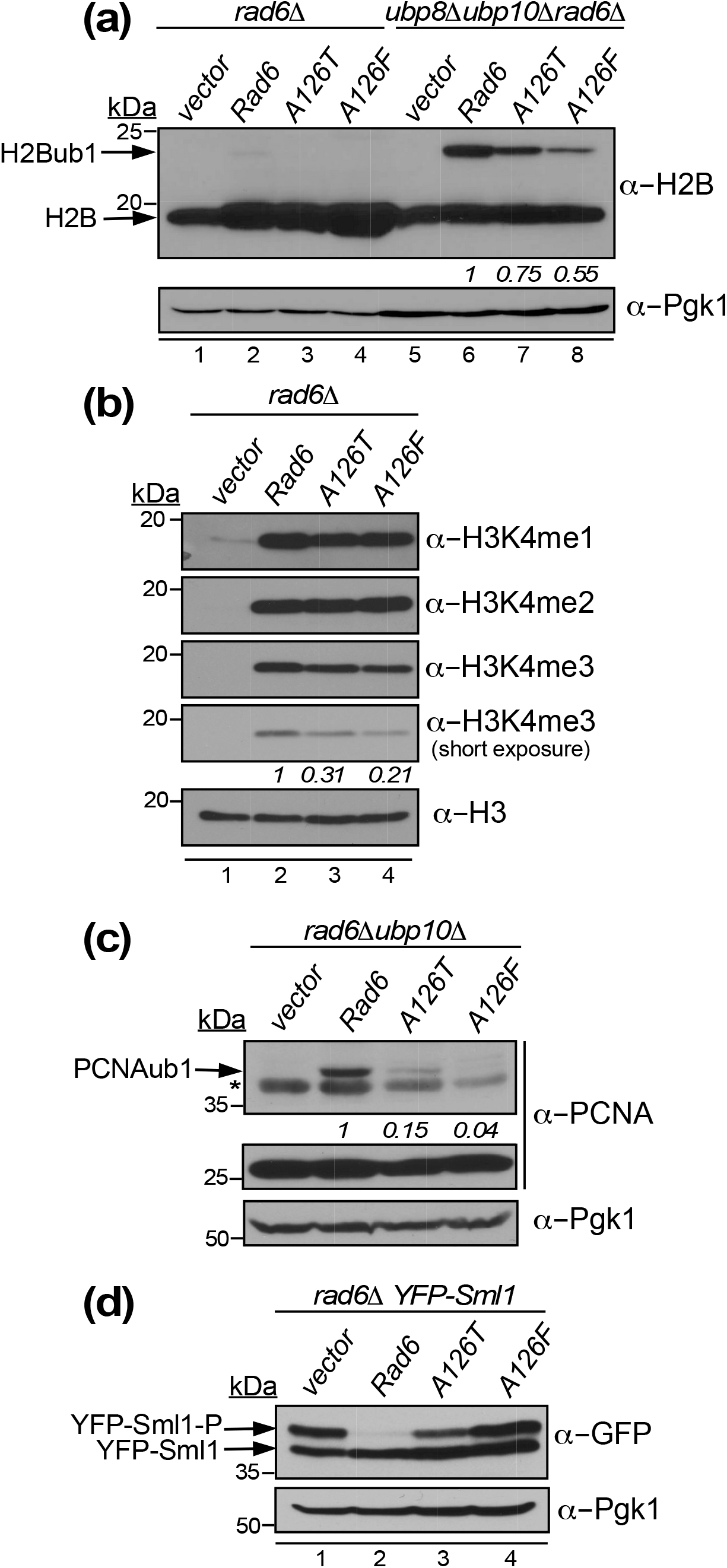
Mutations in A126 of Rad6 impair its target protein ubiquitination functions *in vivo*. (a) Immunoblot for H2Bub1 in a *rad6Δ* strain and in a strain that lacks Rad6, Ubp8 and Ubp10 transformed with empty vector or constructs to express wild-type Rad6 or the mutants rad6-A126T or rad6-A126F. H2Bub1 levels quantified by densitometry relative to strain that expresses wild-type Rad6 are shown for the triple mutant. (b) Immunoblot for histone H3K4 methylation (mono, me1; di, me2; tri, me3) in extracts prepared from the *rad6Δ* strain that expresses the indicated proteins. Histone H3 levels served as loading control. H3K4me3 and H3 levels were quantified by densitometry. H3K4me3 levels normalized to H3 levels in the mutants are shown relative to the strain that expresses wild-type Rad6 (set as 1). (c) Immunoblot for monoubiquitinated PCNA monoubiquitination (PCNAub1) in the *rad6Δ ubp10Δ* strain transformed with empty vector or constructs to express wild-type Rad6 or the indicated mutants and treated with 0.02% methyl methane sulfonate for 90 min. Pgk1 served as loading control. For each strain, PCNAub1 and Pgk1 levels were quantified by densitometry. PCNAub1 levels normalized to Pgk1 levels in a mutant are shown relative to that in a control strain expressing wild-type Rad6 (set as 1). (d) Immunoblots for phosphorylated and unphosphorylated YFP-tagged Sml1 expressed in *rad6Δ* strain and transformed with empty vector or constructs to express wild-type Rad6 or the indicated mutants. Cultures were treated with 3µg/ml bleomycin for 45 min, followed by recovery from DNA damage for 35 min in fresh medium without bleomycin. Pgk1 levels served as loading control. In all panels, molecular weights of the protein standards used as size markers are indicated.

In yeast, steady-state H2Bub1 levels are a net result of two opposing enzymatic activities: Rad6-Bre1-Lge1-mediated ubiquitin conjugation and subsequent removal by deubiquitinases Ubp8 and Ubp10 (60,61). To directly examine the effects of A126 mutations on histone H2B monoubiquitination, we measured the steady-state H2Bub1 levels upon expression of rad6-A126T and rad6-A126F mutants in a *rad6Δ* strain that additionally lacks Ubp8 and Ubp10 (i.e., *ubp8Δ ubp10Δ rad6Δ*). In this background, the rad6-A126T mutant caused a 25% reduction in H2Bub1 levels, and the rad6-A126F mutant caused about 50% decrease when compared to the control strain with wild-type Rad6 (Figure 2a, compare lanes 7-8 to lane 6). These results demonstrate that mutations at A126 compromise the ability of Rad6 to catalyze histone H2B monoubiquitination *in vivo*. Immunoblotting also revealed that the steady-state levels of H3K4me3 were decreased in *rad6-A126T* and *rad6-A126F* mutants when compared to the control strain (Figure 2b), as expected given the reduced H2Bub1 levels and telomeric silencing defects observed in these mutants (Figures 1c and 2a). Moreover, the observed decrease in the H2Bub1-dependent H3K4me3 levels explains the telomeric gene silencing defect of Rad6 A126 mutants.

Following DNA damage, Rad6 and Rad18 E3 ligase catalyze PCNA monoubiquitination (PCNAub1) (28) (Figure 1a). H2BK123 deubiquitinase Ubp10 also removes the ubiquitin from PCNAub1 (62). We therefore examined PCNAub1 levels in the rad6-A126T and rad6-A126F mutants expressed in a *rad6Δ ubp10Δ* double-deletion strain after induction of DNA lesions using methyl methane sulfonate (MMS) treatment. The rad6-A126T and rad6-A126F mutants had severe reductions in the DNA damage-induced PCNAub1 levels when compared to the control strain with wild-type Rad6 (Figure 2c). After DNA damage, Sml1 is phosphorylated and then targeted for proteasomal degradation in a Rad6-Ubr2-Mub1-dependent manner (38). Consistent with this reported study, we observed that the slow-migrating phosphorylated Sml1 was eliminated in the presence of Rad6 during the recovery phase after exposure to DNA damaging agent bleomycin, but it persisted in the absence of Rad6 (Figure 2d, compare lanes 1-2). Phosphorylated Sml1 was retained in both rad6-A126T and rad6-A126F mutant strains upon recovery from DNA damage, (Figure 2d, lanes 3-4), with the latter mutant showing a level of retention similar to that in the *rad6Δ* null strain (Figure 2d, see lanes 1 and 4). Therefore, mutations in A126 compromise the ability of Rad6 to perform mono or polyubiquitination of histone H2B, PCNA, Sml1, and likely other substrate proteins *in vivo*. Thus, providing a molecular explanation for the observed defects in telomeric gene silencing, N-end rule degradation, and DNA repair in yeast cells with mutations in A126 of Rad6.

### A126 mutations do not disrupt interactions of Rad6 with its partner E3 ubiquitin ligases

Ubiquitination of substrate proteins by Rad6 *in vivo* is accomplished via its partnership with different E3 ligases: Ubr1, Ubr2, Bre1, and Rad18 (Figure 1a). To investigate whether defective substrate protein ubiquitination by Rad6 A126 mutants is due to disruption of the association with these partner E3 ligases, we examined the interactions of wild-type or mutant Rad6 with E3 ligases *in vitro* using co-purification and within yeast cells by co-immunoprecipitation (co-IP). Published studies have delineated amino acids 1-214 of Bre1 and 301-487 of Rad18 as the minimal Rad6-binding regions *in vitro* (45-47). Therefore, we co-expressed wild-type or mutant Rad6 tagged with hexahistidine (His_6_) along with these minimal Rad6 binding regions of Bre1 or Rad18 (Bre1R6BR or Rad18R6BR, respectively) in bacteria. Bacterial lysates prepared after co-expression were subjected to metal affinity purification to capture His_6_-tagged wild-type or mutant Rad6 and co-purifying Bre1R6BR or Rad18R6BR, which were then evaluated by SDS-PAGE and Coomassie blue staining. The amounts of Bre1R6BR or Rad18R6BR that co-purified with His_6_-rad6-A126T and His_6_-rad6-A126F were increased or were similar, respectively, to the amounts that co-purified with control His_6_-Rad6 (Supplementary Figure 1a-b).

For co-immunoprecipitation, we expressed Flag epitope-tagged wild-type or mutant Rad6 in a *rad6Δ* strain that also contained Myc-tagged Rad18, V5-tagged Ubr1, or HA-Ubr2, which enable their detection by immunoblotting. Immunoaffinity purification using anti-Flag antibody and subsequent immunoblotting showed that the levels of Ubr1, Ubr2, Rad18, and Bre1 that co-precipitated with rad6-A126T-Flag or rad6-A126F-Flag were similar to or slightly increased to their amounts that co-precipitated with wild-type Rad6-Flag (Supplementary Figure 1c). Taken together, these data from co-purification and co-immunoprecipitation experiments showed that mutations in A126 do not abolish or diminish the interactions of Rad6 with its partner E3 ligases necessary for target protein ubiquitination *in vivo*. Importantly, these results suggested that the A126 mutations in helix-3 could adversely affect the E3 ligase-independent enzymatic activity of Rad6.

### A126 mutations adversely affect activity and stability of Rad6

To test this possibility, we next examined the *in vitro* ubiquitination activities of rad6-A126T and rad6-A126F mutants. Wild-type Rad6, the catalytic-dead mutant rad6-C88A, rad6-A126T and rad6-A126F were expressed in and purified from bacteria. UBE2B or Rad6b, the human homolog of yeast Rad6, was reported to form ubiquitin chains in solution in the absence of an E3 ligase and a target protein(63). However, yeast Rad6 showed significantly compromised *in vitro* intrinsic ubiquitin chain formation activity when compared to its human homologs (Supplementary Figure 2). *In vitro* in the absence of E3 ligases, Rad6 was reported to non-specifically polyubiquitinate histone proteins (49,64). Consistent with these studies, robust polyubiquitination of histone H2B was observed with wild-type Rad6 in our *in vitro* system (Figure 3a, lanes 2-4). Monoubiquitination was decreased, and polyubiquitination was nearly absent in the assay with the rad6-A126T mutant, and neither was detected in the rad6-A126F mutant similar to that in the catalytically inactive rad6-C88A mutant (Figure 3a). Overall, mutations at A126 compromise the enzymatic activity of Rad6 to ubiquitinate substrate protein *in vitro*.

**Fig. 3.**
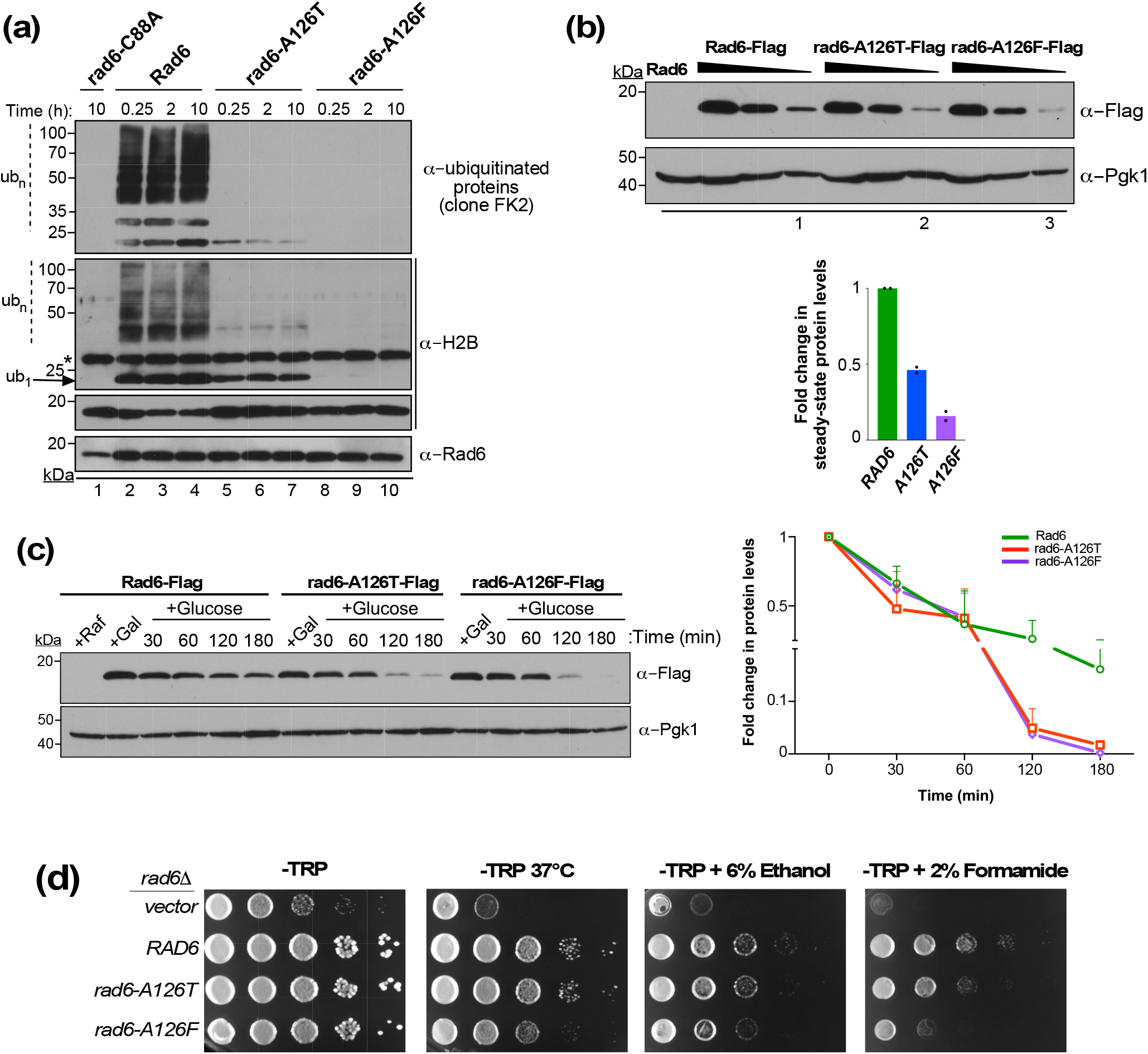
A126 mutations disrupt enzymatic activity and stability of Rad6. (a) Immunoblot of products of an *in vitro* ubiquitination assay with recombinant wild-type Rad6 or indicated mutants. Enzyme was incubated at 30°C for the indicated time along with ubiquitin (Ub), Uba1, ATP/Mg^2+^ and yeast histone H2B (substrate). The reaction mix was then resolved by SDS-PAGE prior to immunoblotting. Ub_1_, monoubiquitinated H2B; Ub_n_, polyubiquitinated H2B. The *asterisk* indicates a cross-reacting protein. The catalytic-dead mutant rad6-C88A served as a control. (b) *Left*, Immunoblot of Flag-tagged wild-type or mutant Rad6. A two-fold serial gradient of the extracts prepared from the indicated strains were resolved by SDS-PAGE prior to immunoblotting. Extract from the strain expressing Rad6 served as a ‘no tag’ control. *Right*, Plot of mean fold-change in the steady-state levels of A126 mutant relative to wild-type Rad6 (± standard error of the mean from two independent experiments) based on densitometry quantitation, for which the signals in the numbered lanes were used. The signals for wild-type or mutant Rad6 were initially normalized to the signals for Pgk1, which serves as a loading control. (c) *Left*, Immunoblot for analysis of stability of Flag epitope-tagged Rad6 or mutants grown in a medium containing raffinose (+Raf; uninduced) or galactose (+Gal; induced), and at different time points after transcription from the *GAL1* promoter was halted by adding glucose. Pgk1 served as loading control and for normalization. *Right*, Plot of mean fold-change in protein levels (± standard error of the mean from two independent experiments) at different time points after glucose-mediated transcriptional inhibition relative to that in the induced state (+Gal, 0 min). (d) Growth assay was conducted by spotting ten-fold serial dilution of the indicated yeast strains on synthetic medium without tryptophan (-TRP) or in medium containing 6% ethanol or 2% formamide and incubated at 30°C. Cells spotted on -TRP medium were also subjected to heat stress by incubation at 37°C.

Next, we examined the effects of A126 mutations on levels of Rad6 *in vivo*. First, whole-cell lysates were prepared from strains expressing Flag epitope-tagged wild-type or mutant Rad6 and subjected to immunoblotting. The global or steady state levels of both rad6-A126T and rad6-A126F mutants were lower than that of wild-type Rad6 (Figure 3c). To determine whether this decrease in the steady state levels was due to compromised protein stability, we expressed a Flag epitope-tagged wild-type or mutant Rad6 in yeast from a galactose-inducible promoter (*GAL1*), and then added glucose to inhibit transcription as previously described (65). Protein levels were then measured at various time points using immunoblotting. The levels of rad6-A126T and rad6-A126F mutants were drastically reduced compared to wild-type Rad6 at 2 hours after glucose-mediated transcription shut-off (Figure 3d), indicating that mutations in A126 are detrimental to the stability of Rad6. Taken together, our results indicate that mutations at A126 compromise both activity and stability of Rad6.

### A126 mutations alter the structure of Rad6

In the reported crystal structure of Rad6 (66), A126 is in close spatial proximity to the catalytic pocket composed of the active-site C88, residues in the HPN motif and the gateway S120 (Figures 1b, Supplementary Figure 3). These residues were shown to be necessary for or to regulate the activity of Rad6 (18,51,67), or E2 enzymes in general (15,22,23). We therefore postulated that the impaired protein ubiquitination or stability of the Rad6 A126 mutants might be due to the adverse effects of these mutations on the overall structure of Rad6.

Growth sensitivity of yeast mutants to high temperature (37°C), ethanol, or formamide indicate general protein structural defects presumably from disrupted hydrogen bonds (68). Spotting assays showed that the rad6-A126T and rad6-A126F mutants have reduced growth at 37°C and in media containing 6% ethanol or 2% formamide when compared to the control strain expressing wild-type Rad6 (Figure 3d), suggesting that mutations at A126 can alter the structure of Rad6.

To test this possibility, we first performed molecular dynamics (MD) simulation, which allows one to evaluate structural details and dynamic behaviors of proteins by measuring the trajectory of individual atoms over time (69-72). *In silico* models of rad6-A126T and rad6-A126F were created using the crystal structure data for native Rad6 (PDB ID: 1AYZ (66)). We then performed all-atom MD simulations for a period of 100 ns. In MD simulations, analysis of root-mean-square deviation (RMSD) of the backbone or Cα atoms provide an overall view of the changes occurring in a protein over the course of the simulation (73). In the RMSD plot, both native Rad6 and the mutants showed very similar patterns of deviations over the timescale of simulation (Supplementary Figure 4a), suggesting that the A126 mutations do not alter the overall structural topology of Rad6.

To determine the impact of mutations on local flexibility and dynamic behavior of individual amino acids, we calculated root-mean-square fluctuation (RMSF) values for the backbone residues in native as well as mutant Rad6 (74-76). Both mutants had higher RMSF values, indicating higher flexibility, than the native Rad6 at residues in the vicinity of T126 or F126 including those in helix-3 and in the adjoining loop-8 region (amino acids 114-S120) (Figure 4a). Additional subtle increases or decreases in RMSF values were also observed for one or both mutants at multiple spatially close and distant residues including those near the catalytic C88 residue. These results suggest that mutation in A126 can either enhance or constrain the flexibility of individual residues to cause local conformational changes in Rad6.

**Fig. 4.**
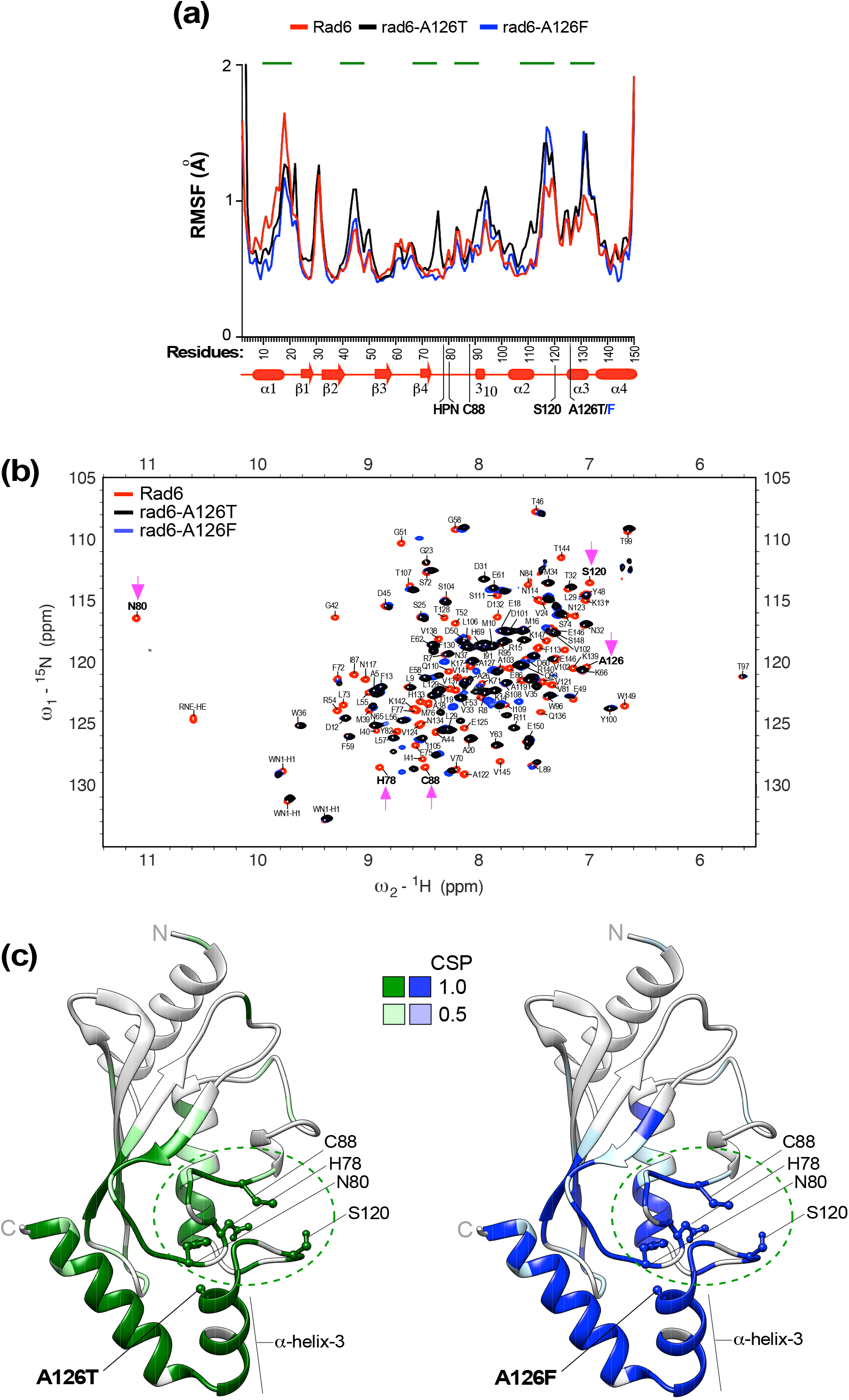
A126 mutations disorganize local as well as global structure of Rad6. (a) RMSF analyses of Rad6 (*red*), rad6-A126T (*black*) and rad6-A126F (*blue*). *Green lines* indicate residues with increased or decreased RMSF values in a mutant relative to wild-type Rad6, indicative of enhanced or constrained flexibility, respectively. Schematic below the x-axis shows the secondary structures of Rad6. The site of mutation in helix-3, the active-site C88, and other catalytically crucial amino acids of Rad6 are indicated. (b) Overlay of the ^1^H-^15^N HSQC spectra of rad6-A126T (*black*) and rad6-A126F (*blue*) mutant on the spectrum of Rad6 (*red*). *Magenta arrows* point to either a complete absence of NMR signal or a drastic chemical shift perturbation for the indicated residues of the catalytic pocket in the mutants compared to wild-type Rad6. (c) Chemical shift perturbations (CSPs) in the A126 mutant mapped onto the Rad6 crystal structure Rad6 (PDB: 1AYZ) using UCSF Chimera. CSPs were quantified for each mutant from the overlay of their NMR spectrum with that of wild-type Rad6 (panel b). No overlap of the NMR signal in the mutant relative to the wild type was scored as 1, and partial overlap was scored as 0.5. Key residues of the catalytic pocket (*green dotted circle*) and the site of each A126 mutation in helix-3 are indicated. *N*, N-terminus; *C*, C-terminus.

To further test this possibility, we performed time-dependent secondary structure fluctuation analysis using DSSP (77), which can yield additional information on the structural flexibility of proteins. The DSSP plot for native Rad6 shows that many of its secondary structures are stable and remain unchanged during the course of MD simulation. However, certain secondary structures, such as helix-3, are flexible and are converted to a 3_10_-helix or turns during the simulation but revert to their original state at the end of the simulation (Supplementary Figure 4b). The DSSP plots for the two mutants show reorganization of the 3_10_-helix (residues 90-93) close to the active-site C88 into a α-helix (Supplementary Figure 4b). Moreover, the helix-2 and helix-3 are reorganized into 3_10_-helix or turns in the rad6-A126F mutant (Supplementary Figure 4b). These findings indicate that mutations in A126 can alter the flexibility and/or conformations of the secondary structures within Rad6.

Given that secondary structure elements in proteins are stabilized by hydrogen bonding, we used the *hbond* tool to analyze the extent of hydrogen bonding in the native and mutant Rad6 proteins (78). The number of hydrogen bonds were reduced during the MD simulation in the rad6-A126F mutant compared to native Rad6, although the stability of rad6-A126T mutant was similar to that of wild-type Rad6 (Supplementary Figure 4c). Computational stability prediction using DynaMut 2.0 (79) further indicated that rad6-A126T and rad6-A126F are both destabilized relative to the native protein (Table 1). Taken together, these computational analyses indicated that mutations in A126 can alter the flexibility of individual residues and/or conformations of local secondary structures of Rad6, in particular those within or near the catalytic pocket. Moreover, these observations match well with the general protein structural defects displayed by the rad6-A126T and rad6-A126F mutants in growth assays (Figure 3d).

**Table 1.** Folding free energy (ΔΔG) and stability predictions using Dynamut 2.0

| Protein | Mutation | $\Delta\Delta G^{\text{stability}}$ (kcal/mol) | Stability change |
| --- | --- | --- | --- |
| Rad6 | A126T | −1.22 | Destabilizing |
|  | A126F | −0.62 | Destabilizing |
| UBE2A | A126T | −1.49 | Destabilizing |
|  | A126F | −0.80 | Destabilizing |
| UBE2B | A126T | −1.83 | Destabilizing |
|  | A126F | −0.73 | Destabilizing |

### A126 mutations perturb the catalytically relevant residues of Rad6

To experimentally validate the data from MD simulations and to directly examine the structural changes caused by the Rad6 A126 mutations, we then employed nuclear magnetic resonance (NMR). We expressed and purified ^15^N isotope-labeled wild-type Rad6 and the mutants rad6-A126T or rad6-A126F from bacteria and recorded their two-dimensional ^15^N-^1^H heteronuclear single quantum coherence (HSQC) NMR spectra. Complete residue assignments were performed on the NMR spectra obtained for wild-type Rad6, and a published dataset was also used (24). The NMR spectrum for each mutant was then overlaid on the assigned NMR spectra for wild-type Rad6 (Figure 4b). NMR chemical shifts are very sensitive to change in protein structure and dynamics; and arise from small differences in the local magnetic field and shifts in equilibria. For a given residue, a perfect overlap of NMR signals between the mutant and wild-type Rad6 indicates no structural perturbation. On the other hand, partial or no overlap in NMR signals between a mutant and wild-type indicates a structural anomaly or a chemical shift perturbation caused by the introduced mutation. The HSQC plot showed no or poor overlap of NMR signals at multiple residues in the rad6-A126T and rad6-A126F mutants when compared with the signals in the wild-type Rad6 spectrum (Figure 4b). Quantitation of chemical shift perturbations showed that A126 mutations alter the positions of residues in the helix-3 and in the adjacent helix-4 (Supplementary Figure 5, Figure 4c). Additional perturbations were also evident at distant sites on the backside of Rad6 that are implicated in non-covalent interactions with ubiquitin (24).

Importantly, the A126 mutations caused perturbations in residues H78, N80, the active-site residue C88 and the gateway residue S120 (Figure 4b-c), which together constitute the catalytic pocket of Rad6. Thus, these NMR data confirmed the findings of MD simulations and demonstrate that mutations in A126 perturb the structure of Rad6, importantly at residues crucial for its enzymatic activity.

Overall, these results from MD simulations and NMR along with the growth phenotypes together demonstrate that mutations in helix-3 cause structural perturbations including at the catalytic pocket, and thus provide an explanation for their adverse effects on enzymatic activity and overall protein stability of Rad6.

### A126 mutations alter the structures of human Rad6 homologs UBE2A and UBE2B

The alanine at position 126 in helix-3 is evolutionarily conserved from yeast to humans (Supplementary Figure 3). Therefore, we then investigated the effects of A126T or A126F mutation on the structures of UBE2A and UBE2B, the human homologs of yeast Rad6. Computational stability prediction (79) indicated that A126T and A126F are both destabilizing mutations in UBE2A and UBE2B (Table 1). Next, we created *in silico* models for threonine or phenylalanine substitution at A126 using the reported crystal structures for UBE2A (PDB ID: 6CYO) (80) and UBE2B (PDB ID: 2YB6) (63) in order to examine their effects on protein structure and conformation using MD simulations. MD simulations for the A126T mutation in UBE2A or UBE2B showed the following: RMSD plots for UBE2A-A126T and UBE2B-A126T mutants showed patterns of deviations very similar to those of their respective native proteins (Supplementary Figure 6a), indicating that threonine substitution at 126 does not cause gross changes in overall topologies of these human proteins. However, higher RMSF values, indicating increased flexibility, were observed for both UBE2A-A126T and UBE2B-A126T at residues near the introduced mutations in helix-3 including in the active-site cleft containing the gateway S120 residue when compared to their respective native proteins (Figure 5a). Consistent with this increased local flexibility, DSSP plots showed disorganization of secondary structures, such as loop-8, which contains the gateway residue S120, and helix-4 in both UBE2A-A126T and UBE2B-A126T mutants at the end of MD simulation (Supplementary Figure 6b). The RMSF plot for the UBE2B-A126T mutant also showed decreased RMSF values indicative of reduced flexibility for residues in the backside loop-3 (amino acids 42-51), β-sheet-3, and helix-4 when compared to native UBE2B (Figure 5a). Moreover, *hbond* analysis showed that the A126T mutation increased the number of hydrogen bonds during the course of the simulations of both UBE2A and UBE2B relative to the native protein (Supplementary Figure 6c), implying that the threonine substitution decreases overall flexibility or causes compaction of the structures of UBE2A and UBE2B.

**Fig. 5.**
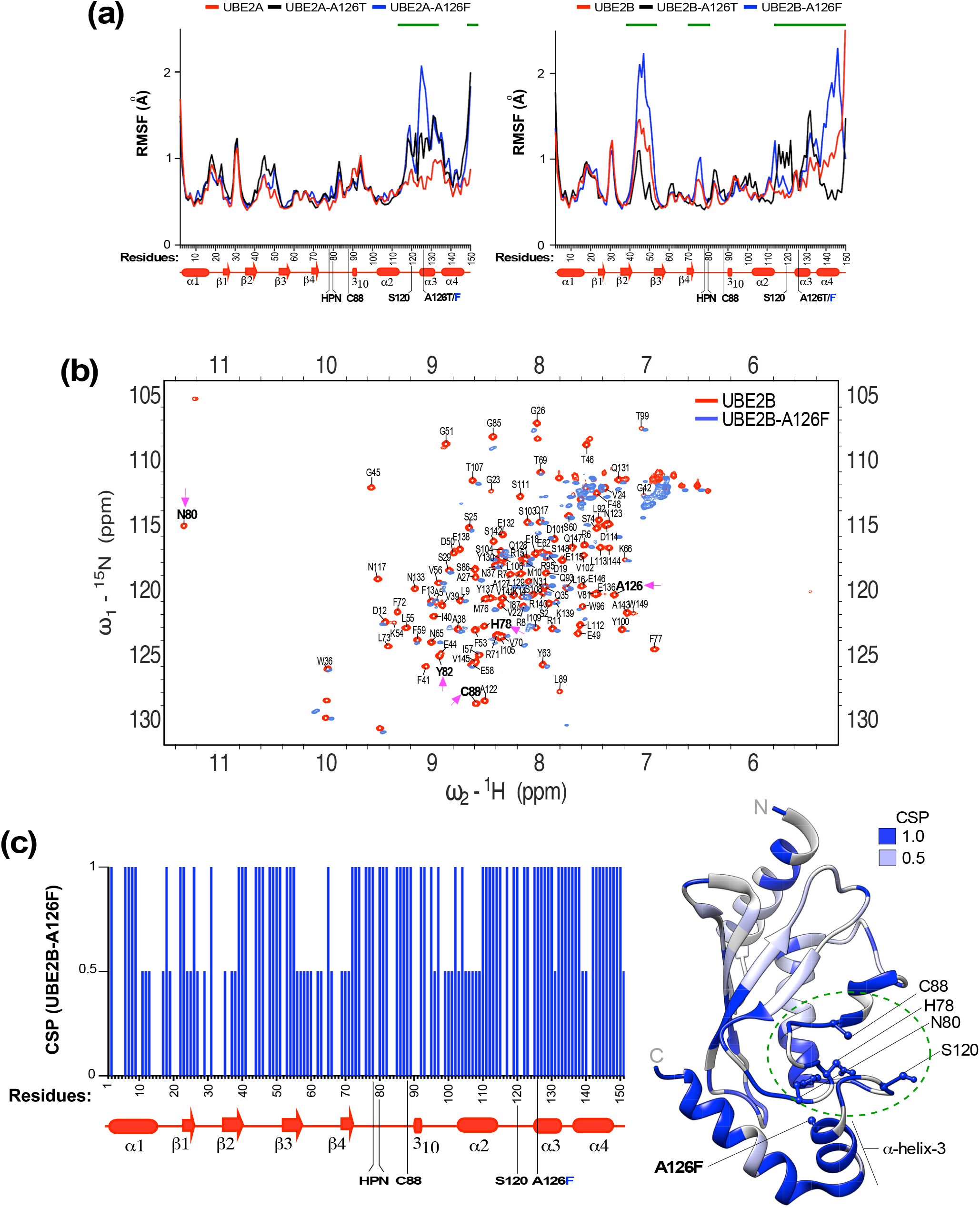
A126 mutations disorganize the structures of UBE2A and UBE2B. (a) RMSF analyses of wild-type UBE2A or UBE2B (*red*) and proteins with mutations A126T (*black*) and A126F (*blue*). *Green lines* indicate residues with increased or decreased RMSF values in the mutants relative to wild-type UBE2A or UBE2B. Schematics below the plots shows the secondary structures of the proteins. (b) Overlay of the ^1^H-^15^N HSQC spectrum for UBE2B-A126F (*blue*) on that obtained for wild-type UBE2B (*red*). *Magenta arrows* point to a complete absence of NMR signal or a drastic chemical shift perturbation for the indicated residues of the catalytic pocket in the mutant compared to wild-type. (c) *Left*, Histogram of the chemical shift perturbation (CSP) at each residue in the mutant. Schematic below the histogram shows the positions of various secondary structures. *Right*, CSPs, scored as described in Figure 4c, mapped onto the crystal structure of UBE2B (PDB: 2YB6) using UCSF Chimera. Key residues of the catalytic pocket of UBE2B (*green dotted circle*) and the site of A126F mutation in helix-3 are indicated. *N*, N-terminus; *C*, C-terminus.

MD simulations for the A126F mutation in UBE2A or UBE2B showed the following: The RMSD plot for the UBE2B-A126F mutant showed significant deviation from that of native UBE2B (Supplementary Figure 6a, top right panel), suggesting that the bulky phenylalanine substitution in helix-3 adversely impacts the backbone Cα atoms to drastically alter the global structure of UBE2B. High RMSF values indicating increased flexibility were evident for residues adjacent to the introduced mutation in helix-3 including the gateway S120 residue in both UBE2A-A126F and UBE2B-A126F mutants when compared to their native proteins (Figure 5a). Moreover, considerable increases in flexibility were also evident in distant-site residues of the backside region in loop-3 and β-sheet-3 in the UBE2B-A126F mutant (Figure 5a). Matching well with the destabilization of local and/or global protein structure, *hbond* analysis revealed that the A126F substitution decreased the number of hydrogen bonds in UBE2B during the course of the MD simulation (Supplementary Figure 6c, right panel). The DSSP plots further accentuated the destabilizing effect of the A126F mutation, as multiple secondary structures in both UBE2A and UBE2B were either disrupted or reorganized during the course of the simulation (Supplementary Figure 6b). These computational simulation studies suggested that like their disruptive effects on yeast Rad6, mutations in A126 of helix-3 also adversely affected the structures of human UBE2A and UBE2B, with UBE2B appearing to be more sensitive to structural perturbations from the A126 mutations.

### A126 mutation perturbs the catalytic pocket residues of human Rad6b/UBE2B

Next, we used NMR to experimentally test the effects of A126 mutation on the structure of a human Rad6 homolog. We focused on the UBE2B-A126F mutation, as our computer simulations indicated that it is a severe destabilizing mutation (Figure 5a, Supplementary Figure 6; Table 1). We expressed and purified from bacteria ^15^N isotope-labeled wild-type UBE2B or the mutant UBE2B-A126F and recorded their two-dimensional ^15^N-^1^H heteronuclear single quantum coherence (HSQC) NMR spectra. We performed residue assignments on the NMR spectra obtained for wild-type UBE2B using a published dataset (63). The NMR spectrum for the UBE2B-A126F mutant was then superimposed on the assigned NMR spectra for wild-type UBE2B. There was no overlap of NMR signals was obtained for many residues in the UBE2B-A126F mutant with the wild-type UBE2B (Figure 5b), indicating that the mutation causes severe structural perturbations within the protein. Quantitation of chemical shift perturbations in the UBE2B-A126F mutant relative to native UBE2B and their subsequent placement on the crystal structure showed that perturbations occurred at multiple residues throughout the mutant protein both close to the site of introduced mutation in helix-3 and at distant sites including at the N-terminus (Figure 5c). Importantly, drastic perturbations were observed for residues H78, N80, C88 and S120 that form the catalytic pocket of UBE2B (Figure 5c).

### A126 mutation adversely affects the solubility and activity of human Rad6b/UBE2B

When heterologous proteins overexpressed in bacteria fail to attain a soluble or native conformation and remain unfolded, they form insoluble protein aggregates termed inclusion bodies (81). Wild-type UBE2B was highly soluble when overexpressed in bacteria, (Figure 6a, see lanes 4-5), indicating that it is a well-folded protein. In stark contrast, a large amount of the overexpressed UBE2B-A126F mutant was detected in insoluble pellet fraction (Figure 6a, see lanes 9-10), suggesting that the protein was misfolded or unfolded. This result is consistent with the computational predictions and results from NMR experiments (Figure 5b-c, Supplementary Figure 6, Table 1), and further demonstrates that the A126F mutation in helix-3 destabilizes or disorganizes the protein structure.

**Fig. 6.**
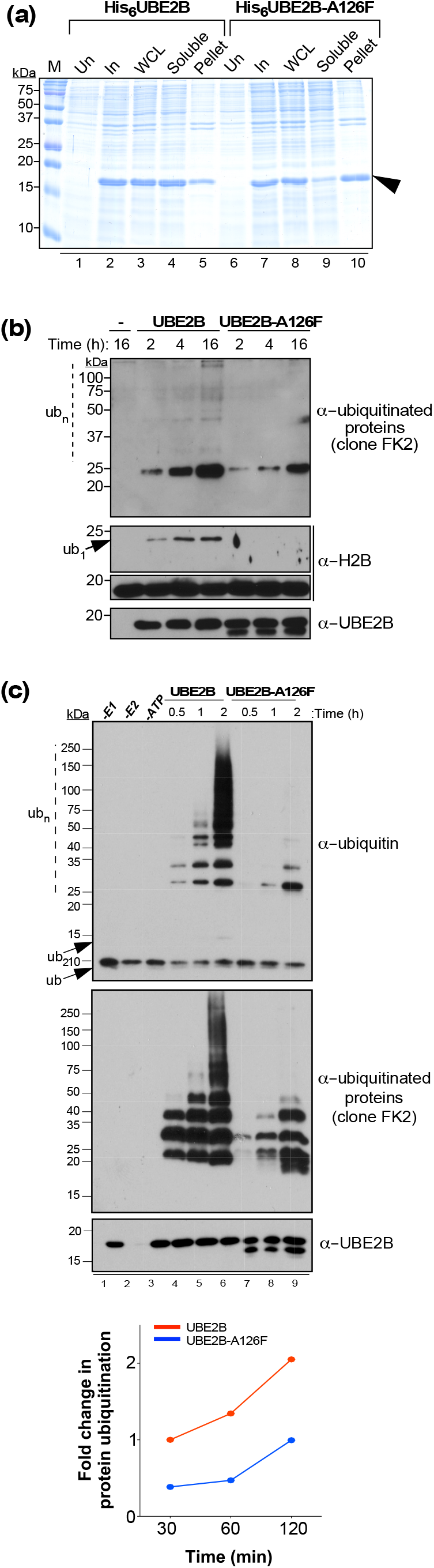
A126 mutation adversely affects solubility and activity of UBE2B. (a) Assessment of solubility of His6-tagged UBE2B or UBE2B-A126F (*arrowhead*) by SDS-PAGE of uninduced (Un) and induced (In) bacterial cells, whole-cell lysate (WCL), and soluble and pellet (or insoluble) fractions. M, protein ladder. (b) *In vitro* ubiquitination assay using recombinant wild-type UBE2B or the UBE2B-A126F mutant was performed essentially as described for Figure 3a; except reactions were incubated at 37°C for the indicated time. Reaction without the E2 enzyme (-) served as a control. *Ub*_*1*_, monoubiquitinated H2B; *Ub*_*n*_, polyubiquitinated H2B. (c) *In vitro* ubiquitin chain formation assay was performed using recombinant wild-type UBE2B or the UBE2B-A126F mutant for the indicated time points. Control reactions without human Uba1 (*-E1*) or UBE2B (*-E2*), or omitting ATP were also performed. Blots were probed with antibodies recognizing ubiquitin, mono- or poly-ubiquitinated proteins (clone FK2) or UBE2B. *Ub*, ubiquitin; *Ub*_*2*_, diubiquitin; *Ub*_*n*_, ubiquitin chains or polyubiquitinated UBE2B. Plot shows quantitation of immunoblots by densitometry. Mean fold-change in the immunoblot signals obtained for the anti-ubiquitinated proteins antibody (clone FK2) for either wild type UBE2B or UBE2B-A126F at indicated incubation times are shown relative to that obtained for wild-type UBE2B after 30 min incubation (set as 1), and were obtained from two independent experiments.

Given the disruptions to the catalytic pocket in the UBE2B-A126F mutant (Figure 5), we performed *in vitro* ubiquitination assays to test the effects of these structural changes on the enzyme activity. Recombinant wild-type UBE2B or the UBE2B-A126F mutant were expressed and purified from bacteria and then used in *in vitro* ubiquitination assays using histone H2B as the substrate. Wild-type UBE2B efficiently mono-or polyubiquitinated the substrate histone H2B, whereas no mono-or polyubiquitinated H2B was observed in the presence of the UBE2B-A126F mutant (Figure 6b). UBE2A and UBE2B show robust *in vitro* ubiquitin chain formation activity in the absence of an E3 ligase or a substrate protein(63) (Supplementary Figure 2). This intrinsic *in vitro* ubiquitin chain formation activity of UBE2B was severely compromised by the A126F mutation (Figure 6c). Collectively, our findings from MD simulations, NMR, and functional assays suggest that mutation in A126 residue in helix-3 disrupts the enzymatic activity of the human Rad6 homolog as well.

## Discussion

Rad6, a multi-functional protein in yeast, regulates telomeric gene silencing via histone H2BK123 monoubiquitination, protein homeostasis by polyubiquitination of N-degron substrates, and DNA repair via monoubiquitination of PCNA and polyubiquitination of Sml1 (28-32,34-36,38-42). Here, we demonstrated that threonine or phenylalanine substitution at A126 in the helix-3 of Rad6 compromises its ability to mono-and polyubiquitinate these target proteins. We also showed that interactions of Rad6 with its various partner E3 ligases, which are necessary for *in vivo* ubiquitination of these target proteins, are not disrupted by mutations at A126. Instead, A126 mutations deform the structure of Rad6 and perturb key residues of the catalytic pocket to inhibit the intrinsic enzymatic activity and to decrease overall protein stability. Thus, our structure-function studies uncover the molecular underpinnings for the phenotypes displayed by yeast with the *rad6-A126T* allele, namely, defects in telomeric silencing, N-end rule degradation and sensitivity to genotoxic agents, which were first reported over two decades ago (53). Moreover, we demonstrated that the bulkier phenylalanine substitution at A126 of helix-3 severely disorganizes local as well as global structures of yeast Rad6 and its human homologs, especially UBE2B, and significantly inhibits their activities. Overall, our studies show that mutations in the conserved helix-3 can disrupt both structure and catalytic functions of yeast and human Rad6 E2 ubiquitin-conjugating enzymes.

Residues in the catalytic pocket of E2 enzymes are implicated in deprotonating the ε- amino group of substrate lysine and converting it into a nucleophile, which then attacks the thioester adduct formed between the E2 active-site cysteine and ubiquitin (Ub) or ubiquitin-like modifications (Ubls) (e.g., SUMO) (82,83). Two mechanisms proposed to explain how residues in the E2 catalytic pocket perform lysine deprotonation are: 1) they may act as proton acceptors for the incoming substrate lysine, as reported for H94 in UBE2G2 and D117 in UBE2D1 (84,85); or 2) they may form a microenvironment that reduces the pKa of the incoming lysine, as reported for residues N85, Y87 and D127 of UBC9, the SUMO-specific E2(82). From the crystal structure of UBC9-SUMO-substrate RanGAP1(86) (Figure 7a), it is evident that the active-site cysteine and the key residues implicated in lysine deprotonation (N85, Y87 and D127) are all in close proximity to the incoming lysine (∼3–5 Å). The optimal distance between the E2 cysteine and the acceptor lysine of the substrate proteins for the transfer of Ub/Ubl is expected to be between 2 – 2.5 Å (87). Therefore, even small perturbations to the conformation of the active-site cysteine and/or other residues of the catalytic pocket of E2 enzymes can alter their operational distance from Ub/Ubl or the substrate lysine to disrupt the conjugation activity.

**Fig. 7.**
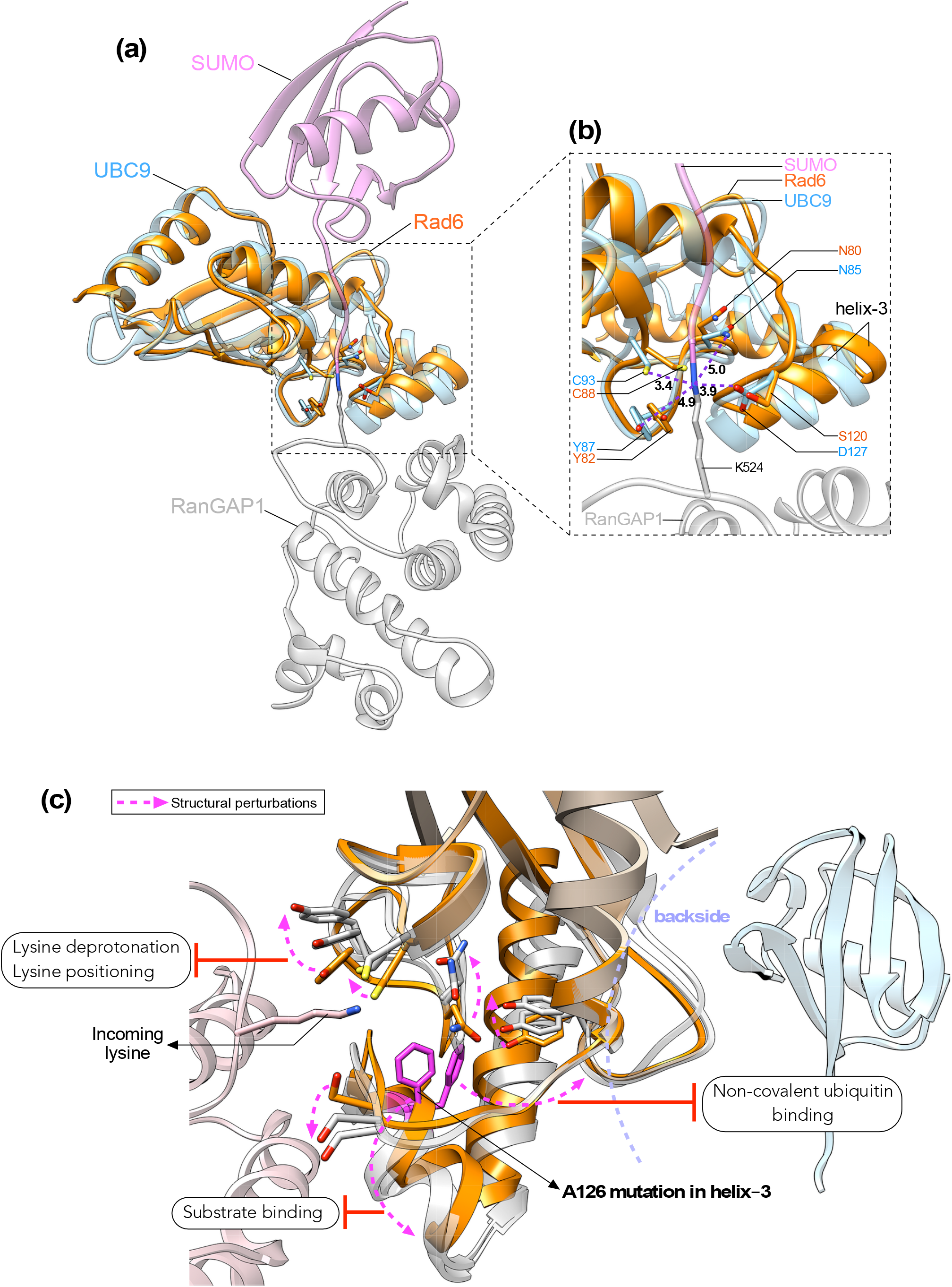
Models for the contributions of helix-3 to the structure and functions of E2 enzymes. (a) Structure of Rad6 (PDB ID: 1AYZ) was superposed onto that of UBC9 E2 enzyme in the co-crystal structure of UBC9-SUMO-RanGAP1 (PDB ID:1Z5S) to show the proximity of helix-3 of Rad6 to its catalytic pocket, ubiquitin, and the incoming lysine of a substrate protein. (b) Zoomed image shows the location and distances of the key residues of the catalytic pocket from the isopeptide bond. Also, shown are the distances in angstroms (Å) of the residues of the catalytic pocket in UBC9 to the isopeptide bond between SUMO and target K524 in substrate RanGAP1. (c) A generalized model to explain how mutation in helix-3 causes local and long-distance structural perturbations (*dotted arrows*) at catalytically key residues and secondary structures of Rad6 or its homologs UBE2A or UBE2B, and perhaps E2 enzymes in general, to inhibit their catalysis-related transactions with ubiquitin or ubiquitin-like modification and substrate proteins.

To address how structural perturbations caused by mutations in helix-3 might impinge on the enzymatic activities of Rad6 and its human homologs, we superposed the structures of yeast Rad6 and human UBE2A or UBE2B onto the reported structure for the Ubc9 E2 in the act of conjugating SUMO onto substrate RanGAP1(86) (Figures 7b; Supplementary Figure 7), which shows that the catalytic pocket residues of yeast and human Rad6 proteins are spatially positioned very similar to their counterparts in UBC9. Importantly, these catalytically key residues are present adjacent to or contiguous with helix-3 in Rad6 and its human homologs. Our MD simulations and NMR experiments show that mutations in helix-3 perturb the active-site cysteine and other residues of the catalytic pockets of yeast and human Rad6 proteins (Figures 4-5). We therefore propose that mutations in helix-3 cause structural changes that alter the distances of the key residues of the catalytic pockets of Rad6 and its human homologs from ubiquitin and/or substrate lysine (Figure 7c). Thus, mutations in the conserved helix-3 can block ubiquitination activity, as demonstrated by our *in vitro* and/or *in vivo* studies, by causing conformational changes to the critical residues involved in catalysis.

In addition to directly altering the position of the active-site cysteine, mutations in helix-3 can also disrupt the ubiquitin-conjugation activity of yeast or human Rad6 enzymes in other ways. In Rad6, UBE2A, UBE2B and other E2 enzymes, the catalytic pocket is buttressed by loop-8 or the active-site cleft, which serves as the gate into the active-site. Studies of UBE2K and UBC13 have shown that the opening and closing of the active-site gate is precisely balanced and that even small deviations in gating impair the functions of UBC13 during DNA damage (88-90). A conserved serine or aspartate, termed the gateway residue, is present in the active-site cleft of E2 enzymes, and regulates ubiquitination (22). S120 is the gateway residue in Rad6 and its human homologs. This amino acid corresponds to D117 of UBE2D1 and D127 of UBC9, which are implicated in deprotonating the incoming substrate lysine (82,84). Phosphorylation of S120 regulates the activities of both Rad6 and UBE2A (51,91). Although S120 as such as cannot act as a proton acceptor, modification of S120 with a negatively charged phosphate can mimic the acidic nature of an aspartate, allowing this residue to act as a proton acceptor or as a pKa reducer for the incoming lysine. Therefore, mutations in helix-3 could sway the gating dynamics to promote either an inactive or a constitutively active conformation or, alternatively, could impact S120 phosphorylation, to affect substrate lysine deprotonation during ubiquitination.

Residues N85 and Y87 of UBC9 are also implicated in reducing the pKa of the incoming lysine (82). These residues correspond to residues N80 and Y82, respectively, of Rad6 and its human homologs. N80 is part of the evolutionarily conserved HPN motif of E2 enzymes. The HPN motif of E2 enzymes functions in localizing the target lysine and in stabilizing the oxyanion formed in the reaction intermediate during the nucleophilic attack (19,21). The asparagine in this motif aids in the formation of the isopeptide bond, histidine is necessary for the structure, and proline promotes the stable transition of these two amino acids (20,21,52,89). Recently, UBE2A-Q93E was reported to be a novel pathogenic mutation associated with mild intellectual disability; and this mutation was proposed to disturb the catalytic microenvironment of UBE2A essential for its substrate lysine deprotonation (80). Q93 of Rad6, UBE2A and UBE2B correspond to the proton acceptor H94 in UBE2G2 (85). Our NMR analyses showed that H78, N80, Y82, and Q93 are all significantly perturbed in rad6-A126T, rad6-A126F and UBE2B-A126F (Figures 4-5). Thus, it is conceivable that mutations in helix-3 can adversely affect the optimal spatial locations of these catalytically vital residues and thus inhibit their functions in substrate lysine deprotonation or oxyanion stabilization during ubiquitination by Rad6 or UBE2A/B (Figure 7c).

E1 ubiquitin-activating enzymes transfer ubiquitin onto the active site cysteine of an E2 enzyme to convert it into an enzymatically active state. Structure-function studies of E2s in complex with Uba1 E1 enzyme have shown that residues of helix-3 are part of the E1-E2 interaction interface and are important for the E1-mediated ubiquitin charging of an E2 enzyme(92-94). Thus, one could envisage mutations in A126 to impede the initial ubiquitin charging step to adversely affect the enzymatic activities of yeast or human Rad6. The backside regions of E2 enzymes, comprised of residues of the four β-sheets, the intervening loops, and the C-terminal ends of helices 1 and 4, is the site of non-covalent interactions with ubiquitin (15,23,24,95). This weak affinity interaction promotes increased processivity of poly-ubiquitin chain formation by E2 enzymes (95). Our RMSF analyses and NMR studies revealed that A126 mutations in helix-3 perturb the conformations of the backside regions of yeast Rad6 and its human homolog(s), especially loop-3 and the C-terminal end of helix-4 (Figures 4-5). Therefore, one could further speculate that the weak-affinity interactions of ubiquitin with the backsides of Rad6 and its human homologs may be abolished because of the structural disruptions caused by helix-3 mutations, which in turn inhibits their polyubiquitination activities, as seen in our *in vitro* or *in vivo* experiments (Figures 1d, 2d, 3a, 6b-c). The proximity of helix-3 to the catalytic pocket and its surface accessibility suggest that it might also play a role in substrate recognition by E2 enzymes. Indeed, the helix-3 of UBC9 interacts with the substrate Ran-GAP1 (82) (Figure 7a). Thus, mutations in helix-3 could prevent substrate binding or correct positioning of the incoming lysine (Figure 7c). In summary, in a simpler analogy, we envision that the conserved helix-3 acts like a lower jaw that controls the movements or functioning of the lips, which correspond to the catalytic pocket of the E2 enzymes.

### Experimental procedures

#### Yeast strains and media

Yeast cells were grown in YPAD broth (1% yeast extract, 2% peptone, 2% dextrose, and 0.004% adenine hemisulfate) or in synthetic dropout (SD) media. Agar (2%) was added to liquid broth to prepare solid media. To create gene knockout strains, the coding region of a target gene was replaced in the parental strains (YMH171 (96) and/or DHY214/DHY217) or the W4622-14B (38) strain using PCR products containing ∼500 bp each of the promoter and terminator regions of the target gene and the open reading frame (ORF) replacement *KanMX6* selection cassette, which were amplified using genomic DNA isolated from the respective deletion mutant strain from the Open Biosystem’s yeast deletion collection. Alternatively, a one-step PCR-based gene knockout strategy was performed using pF6a-KanMX or pAG25 (natMX4) or pAG32 (hphMX4) (97) as the template. The *RAD6* coding region was replaced with *URA3* using a construct that contained the *RAD6* promoter and terminator sequences flanking the *URA3* gene, which was linearized with HindIII-BamH1 prior to transformation. YMC309 and YMC336 double or triple gene knockout strains were created by mating of single or double gene deletion strains, followed by sporulation and tetrad dissection. Genotypes of yeast strains used in this study are listed in Table 2.

**Table 2.**
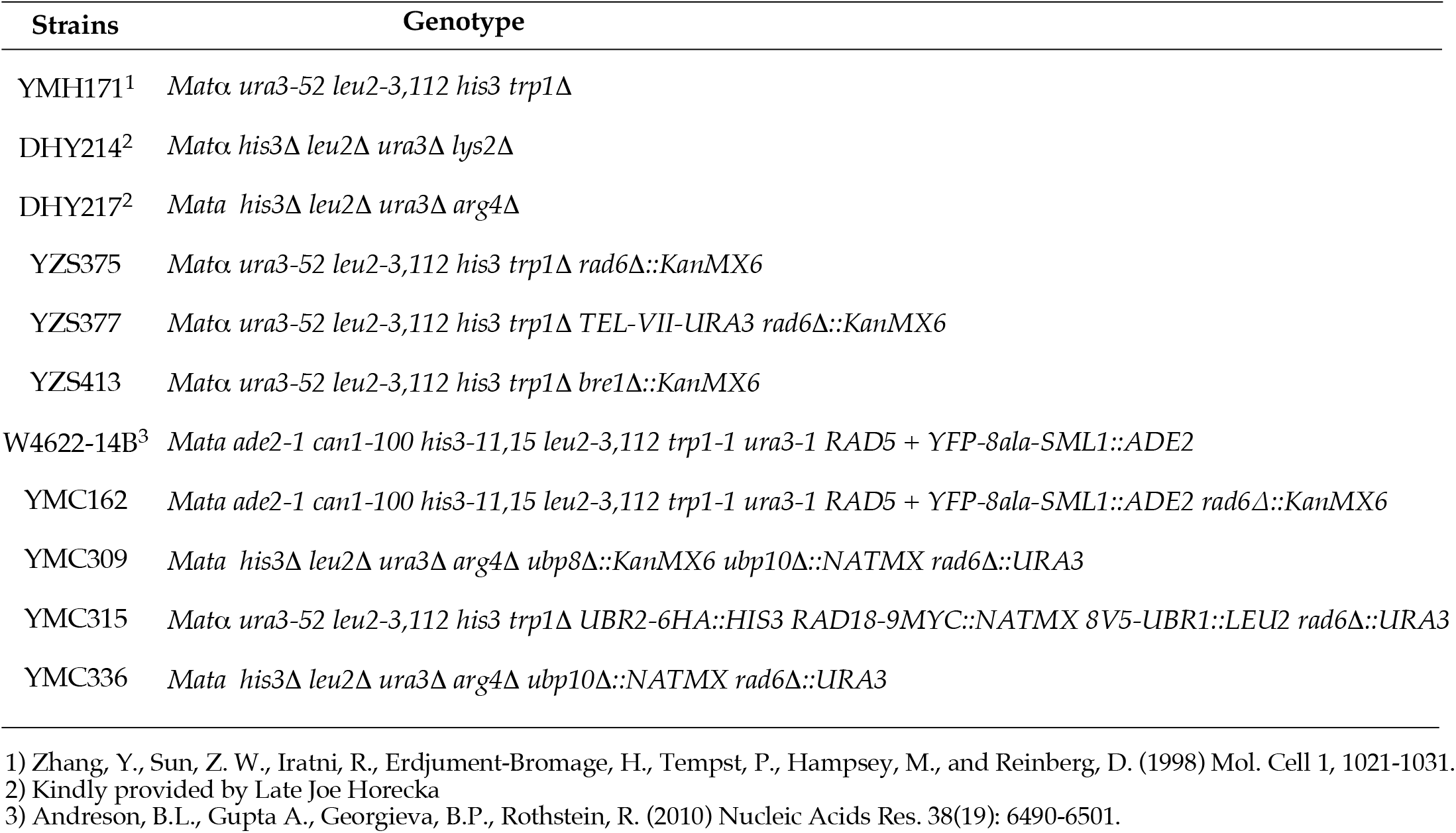
Yeast strains list.

#### Plasmid constructs

The *RAD6* terminator region (450 bp) was PCR amplified using yeast genomic DNA as the template. The PCR product also contained sequences for a Flag epitope-tag, a stop codon, Spe1 and BamH1 sites at the 5’-end and a Kpn1 site at the 3’-end. This PCR product was digested with Spe1-Kpn1 and inserted into the same sites in vector pRS314 (*TRP1*, CEN). Into this construct, the *RAD6* promoter region (286 bp), PCR amplified using yeast genomic DNA as template was inserted as a Not1-Spe1-digested fragment, to obtain construct pMC5 (*RAD6 promoter-Spe1-BamH1-Flag-RAD6 terminator, TRP1* CEN). The coding sequence for wild-type Rad6 without a stop codon was PCR amplified using yeast genomic DNA as the template and additionally contained Spe1 and BamH1 sites at its 5’ and 3’ ends, respectively. This PCR product and construct pMC5 were digested with Spe1 and BamH1 prior to their ligation using T4 DNA ligase (Invitrogen). Substitution mutations were introduced into Rad6 using a PCR-based site-directed mutagenesis approach. For galactose-inducible expression, the DNA fragments encoding Flag-tagged Rad6, rad6-A126T, or rad6-A126F and the terminator sequence were excised using Spe1 and Kpn1 and inserted into the same sites in vector pRS316 (*GAL1 URA3*, CEN).

For proteins used in NMR studies, the wild type or mutant *RAD6* coding region (amino acids 1 – 150) was PCR amplified from yeast constructs described above and inserted into the Nde1 and BamH1 sites downstream of sequence encoding the His_6_ tag and the thrombin cleavage sequence in bacterial expression vector pET28a (Novagen). IDT g-blocks® fragments were synthesized for wild-type UBE2A or UBE2B or their mutants and inserted into Nde1-BamH1-digested pET28a by sequence and ligation independent cloning (SLIC) (98). For co-expression of Rad6 (or its mutants) and Bre1, the coding region (amino acids 1 – 150) of wild-type Rad6 or an A126 mutant was PCR amplified and inserted by SLIC into BamH1-Not1 sites in bacterial expression vector pRSF-Duet (Novagen). Subsequently, the sequence encoding the Bre1 Rad6-binding region (R6BR) (amino acids 1 – 214) was PCR amplified and inserted between BglII-Xho1 sites using SLIC. For co-expression of Rad6 (or its mutants) and Rad18, the sequence encoding the Rad18 Rad6-binding region (R6BR) (amino acids 301 – 487) and that of the coding region of glutathione-S-transferase (GST) were PCR amplified and then inserted into Nde1-Xho1-digested pRSF-Duet constructs containing wild-type Rad6 or an A126 mutant sequences using SLIC. The GST protein tag ensured solubility of Rad18R6BR when expressed in *E. coli*.

#### Spotting assays

Telomeric silencing reporter strain YZS377 was transformed with either vector pRS314 (*TRP1*, CEN) (99) alone or construct pMC5 or derivatives containing either wild type *RAD6* or a mutant (A126T or A126F). These strains were grown overnight at 30°C with constant shaking in liquid SD media lacking tryptophan (-TRP). Cells (1 OD_600_ or 1 × 10^7^) were harvested, and a 10-fold serial dilution was performed prior to spotting them onto solid -TRP media. For the silencing assay, the media additionally contained 5-fluroorotic acid (5-FOA) and cells were grown at 30°C for 2-3 days. For UV and other drug sensitivity assays, strain YZS375 was transformed with a plasmid construct to express either wild-type Rad6 or one of the mutants (rad6-A126T or rad6-A126F). For the UV sensitivity assay, cells were grown and serially diluted as described above and spotted onto solid -TRP plates prior to being exposed to 254 nm UV light for 15 s. The plates were incubated for 2 days at 30°C prior to imaging. For drug or compound sensitivity assays, serially diluted cells were spotted onto -TRP media 4% bleomycin, 6% ethanol or 2% formamide. For examining heat sensitivity, cells spotted on -TRP plates were incubated for 2 days at 37°C.

#### β-galactosidase assay

Yeast strain (YZS375) was transformed with either vector pRS314 (*TRP1*, CEN) or construct pMC5 derivative containing either wild type *RAD6* or a mutant (*rad6-A126T* or *rad6-A126F*) along with an N-end rule reporter plasmid (pUB23-R-betagal *URA3*, 2μ)(57). These strains were grown in SD media without tryptophan and uracil (SD-TRP-URA), and with raffinose as the sugar source. Reporter expression was induced by the addition of 2% galactose. The LacZ assay was performed using the Yeast β-Galactosidase Assay Kit (Thermo Scientific) and by following the manufacturer’s microfuge tube protocol. Three technical and biological replicates were performed for each strain.

#### Protein expression

The pET-28a-based constructs for the expression of His_6_-tagged wild-type or mutant Rad6 or UBE2B were transformed into *E. coli* strain BL21-CodonPlus(DE3)-RIL (Agilent Technologies). For expression and purification of the human E1 enzyme, pET3a-hUBA1 (Addgene#63571, kindly provided by Dr. Titia Sixma)(100) was transformed into *E. coli* strain BL21-CodonPlus(DE3)-RIL. Overnight cultures were used to seed fresh 1 L of Luria broth (LB) medium containing 50 μg/ml kanamycin and 10 μg/ml chloramphenicol at OD_600_ 0.1 and then grown with shaking at 37 °C to an OD_600_ 0.6. Protein expression was induced by adding 0.5 mM IPTG (GoldBio) and grown overnight at 16°C. Cells were harvested by centrifugation 6000 rpm for 15 min at 4°C. The expression plasmid for yeast H2B in pET11a was transformed into *E. coli* BL21-CodonPlus(DE3)-RIL strain and overnight cultures were used to seed 1L of LB medium containing 100 μg/ml ampicillin and 10 μg/ml chloramphenicol at OD_600_ 0.1, and then grown with shaking at 37 °C to OD_600_ 0.6. Protein expression was induced by adding 1 mM IPTG (GoldBio) and grown at 37°C for 5 h with agitation.

All isotopically labeled proteins were produced in *E. coli* BL21-CodonPlus (DE3)-RIL (Agilent Technologies). To generate the isotopically labeled Rad6, expression was carried out in M9 minimal media supplemented with 3 g/L (^13^C_6_, 99%)-D-glucose and/or 1 g/l (^15^N, 99%)-NH_4_Cl (Cambridge Isotope Laboratories). To generate isotopically labeled rad6-A126T, rad6-A126F, UBE2B or UBE2B-A126F, the M9 minimal media was supplemented with 1 g/L (^15^N, 99%)-NH_4_Cl and 10 g/L D-glucose. Bacterial cultures were grown in M9 minimal media containing 50 μg/ml kanamycin and 10 μg/ml chloramphenicol at 37°C to OD600 ∼ 0.6. Heterologous protein expression was induced with 0.5 mM IPTG (GoldBio), and cultures were grown overnight at 19°C with gentle agitation.

#### Metal affinity copurification

The pRSF duet-based constructs to co-express either wild-type or mutant Rad6 and Bre1R6BR or Rad18R6BR described above were transformed into *E. coli* BL21-CodonPlus(DE3)-RIL strain. Primary cultures (10 ml) were grown overnight at 37 °C in LB medium containing 30 µg/ml kanamycin and 10 µg/ml chloramphenicol. The overnight culture was used to then seed a fresh 10 ml of LB medium with the indicated antibiotics at OD_600_ 0.1 and grown with shaking at 37 °C to OD_600_ 0.6. Protein expression was induced by adding 0.5 mM IPTG (GoldBio) followed by growth overnight at 16°C. Cells were harvested by centrifugation and resuspended in 1mL of lysis buffer (25 mM Tris.Cl pH 7.9, 150 mM NaCl, 20 mM imidazole, 0.1% Triton X-100 and 0.1mM PMSF). The cells were lysed by sonication for 2 min using a Misonix Sonifier and the lysate was clarified by centrifugation at 12000 rpm for 15 min at 4°C. Aliquots of cells or lysates were set aside, pre or post IPTG addition or clarification by centrifugation, to serve as uninduced or induced and whole cell lysates or soluble fractions.

His_6_-tagged proteins in the clarified supernatant (1 ml) were allowed to bind pre-equilibrated TALON^®^ SuperFlow™ resin (Cytiva) with end-over-end mixing for 2 h at 4 °C. Beads were collected by centrifugation at 2,800 rpm for 3 min and then washed three times with lysis buffer. The beads were boiled in 2x Laemmli Sample Buffer (Bio-rad). His_6_-tagged wild-type or mutant Rad6 and co-purifying Bre1R6BR or Rad18R6BR in the eluates were separated by 12% SDS-PAGE and visualized by staining with SimplyBlue™ Safe Stain (Invitrogen). Two independent pull-down or copurification experiments were performed. Stained gels were destained extensively in water, and protein bands were quantified using densitometry (Image J).

#### Protein purification

Whole cell lysates were prepared in Lysis Buffer (25 mM Tris.Cl pH 7.9, 150 mM NaCl, 20 mM imidazole, 1 mM TCEP, and 1 mM PMSF), and were digested with lysozyme (Sigma) for 20 min on ice and then sonicated using a Misonix Sonifier. The soluble fraction was then obtained by high-speed centrifugation (40,000 rpm, 30 min at 4°C) using Ti45 rotor in a Beckman Optima™L90-K Ultracentrifuge. Protein purification from the soluble lysate was performed using a three-step chromatography in an ÄKTA FPLC system (Cytiva). The soluble supernatant was first loaded onto a nickel affinity column (HisTrap™ FF, Cytiva), washed extensively with Lysis Buffer (10 column volumes), and eluted with a 20-500 mM imidazole gradient. Fractions with purified protein were combined, and thrombin (10 units; Sigma) was added to remove the His_6_ tag, except for proteins in *in vitro* ubiquitin chain formation assay. Samples were dialyzed overnight at 4°C into a buffer containing 25 mM Tris.Cl pH 7.9, 50 mM KCl, 10% glycerol, 1 mM EDTA, and 1 mM DTT. The dialysate was centrifuged (40,000 rpm, 30 min at 4°C) and then loaded onto Mono Q anion exchange column (Cytiva) and eluted using a 50-1000 mM KCl gradient. Fractions with purified protein were then loaded onto a Superdex™ 75 gel filtration column (Cytiva) in a buffer containing 25 mM Tris.Cl pH 7.9, 150 mM NaCl, 10% glycerol, 1 mM EDTA and 1 mM TCEP. For NMR, size-exclusion chromatography was performed in a buffer containing 25 mM sodium phosphate buffer pH 7.5 and 0.5 mM TCEP. Eluted fractions in all chromatography steps were analyzed by SDS-PAGE. The final purified proteins were concentrated using a Vivaspin 10 kDa MWCO Centrifugal Concentrator. For NMR, isotopically-labeled and purified UBE2B or UBE2B-A126F were dialyzed into NMR sample buffer (50 mM sodium phosphate buffer pH 8.0, 300 mM NaCl, 1 mM DTT, 10% D_2_O). To purify yeast histone H2B, cell pellets after IPTG induction were resuspended in lysis buffer (50 mM Tris.Cl, pH 7.5, 100 mM NaCl, 1 mM EDTA, and 1 mM PMSF). Cells were lysed by sonication and the lysate was clarified by centrifugation at 40,000 RPM for 30 min in a Beckman Ultracentrifuge. The soluble fraction was discarded, and the pellet (or inclusion body) fraction was dissolved in an unfolding buffer (7 M guanidium chloride, 20 mM Tris.C1 pH 7.5, and 10 mM DTT). Following centrifugation, the supernatant was directly dialyzed first against 1 L SAU-200 (7 M urea, 20 mM sodium acetate pH 5.2, 200 mM NaCl, 5 mM β-mercaptoethanol, and 1 mM EDTA) for ∼6 h in a cold room and then overnight against fresh 1 L SAU-200, also in the cold room. The dialysate was subsequently loaded onto a SP Sepharose FF column (Cytiva) and eluted with SAU-600 (7 M urea, 600 mM NaCl, 20 mM sodium acetate pH 5.2, 5 mM β-mercaptoethanol, and 1 mM EDTA). Fractions containing histone H2B were pooled and dialyzed into water. Refolding of histone H2B was done by dialysis against 2 L refolding buffer (2M NaC1, 10 mM Tris.C1 pH 7.5, 1 mM EDTA, and 5 mM β- mercaptoethanol) overnight at 4°C. Protein in the dialysate was concentrated and purified over a Superdex 75 column in the refolding buffer. Fractions were analyzed on 12% SDS PAGE, and those containing histone H2B were pooled and concentrated using a 10-kDa MWCO Centrifugal Concentrator. For the *in vitro* ubiquitination assay, the purified histone H2B was diluted into a buffer containing 25 mM Tris.C1 pH 7.5, 100 mM NaCl, 1 mM EDTA, and 1 mM DTT.

#### Solubility assay

Overnight cultures for *E. coli* cells expressing recombinant UBE2B or UBE2B-A126F were used to seed 10 ml of LB medium containing 50 μg/ml kanamycin and 10 μg/ml chloramphenicol at OD_600_ 0.1 and grown further at 37°C to OD_600_ 0.6, then induced with 0.5 mM IPTG and grown overnight at 19°C. Cells were harvested and resuspended in a lysis buffer containing 25 mM Tris.Cl pH 7.9, 1 M NaCl, 0.1 mM EDTA, 5 mM imidazole, 1mM TCEP, and 1 mM PMSF. After sonication, the lysate was centrifuged at 10,000 rpm for 15 min at 4°C. Aliquots of the lysate before and after centrifugation were designated as whole-cell lysate and soluble lysate fractions, respectively. The pellet or insoluble fraction obtained after centrifugation was dissolved in 0.5 mL lysis buffer. The samples were then resolved by 15% SDS-PAGE and proteins were visualized by staining using SimplyBlue™ Safe Stain (Invitrogen).

#### *In vitro* ubiquitination assay

The ubiquitination reaction contained 1X Reaction buffer and 5mM Mg-ATP (Ubiquitylation Assay Kit; Abcam), 0.1 µM recombinant yeast or human GST-Uba1/UBE1 (E1, R&D Systems), 2.5 µM recombinant yeast or human ubiquitin (R&D Systems), 0.1 µM wild-type or mutant Rad6 or UBE2B (E2), and 2 µM substrate recombinant yeast histone H2B. For yeast Rad6 or its mutant derivatives, the reaction was performed at 30°C for 15 min, 2h or 10h. For UBE2B or UBE2B-A126F, the reaction was performed at 37°C for 2 h, 4 h or 16 h. The reactions were stopped by adding 2X Laemmli sample buffer (Bio-Rad) and resolved in a 12% SDS-PAGE prior to immunoblotting with anti-mono- and poly-ubiquitinated protein antibody (clone FK2), anti-yeast H2B antibody or anti-Rad6 or UBE2B antibody (see details below).

Ubiquitin chain formation assays were performed essentially as described (63): purified Rad6, UBE2A, UBE2B or their mutants (3 µM) along with 12 µM yeast or human ubiquitin and 90 nM yeast or human E1 enzyme were included in a buffer (50 mM Tris.Cl pH 8.0, 50 mM NaCl, 50 mM KCl, 10 mM MgCl_2_, 0.1 M DTT, 3 mM ATP) and incubated at 30°C (for yeast proteins) or 31°C (for human proteins) at various time points as indicated in the figures. Control reactions lacking either E1 or E2 enzyme or ATP were also performed. The reactions were denatured in SDS-PAGE sample buffer and resolved in Novex™ 4-20% Tris-Glycine gels (Invitrogen) before immunoblotting with anti-ubiquitin or anti-ubiquitinated protein (clone FK2) antibodies or anti-Rad6 or anti-UBE2A or anti-UBE2B antibody (see below). *In vitro* substrate ubiquitination and ubiquitin chain formation assays were confirmed using two independent protein preparations. At least two independent assays were performed and quantitation of immunoblots for anti-ubiquitinated proteins (clone FK2) antibody was performed by densitometry using ImageJ software.

#### Co-immunoprecipitation

Log-phase cultures of yeast cells (50 × 10^7^) expressing Flag epitope-tagged Rad6 or its mutants were harvested, washed once with PBS and stored at -80°C. Cells were lysed by bead beating after resuspension in IP-Lysis Buffer (10 mM Tris.Cl pH 8.0, 100 mM NaCl, 10% glycerol, 0.1% NP-40, and Roche cOmplete™ EDTA-free protease inhibitor cocktail). The samples were cooled on ice for 5 min between the bead-beat cycles and clarified by two high-speed centrifugations (13200 rpm at 4°C) for 20 min and 10 min to obtain the final soluble lysate. Protein estimation was performed using Bio-Rad Protein Assay. An aliquot of the whole cell lysate (50 μg) was set aside for ‘input’. Lysate (1 mg) from various yeast strains was used in immunoprecipitation with anti-Flag M2 magnetic beads (20 μl, Sigma) in a total volume of 1.5 ml of IP-Lysis buffer and incubated with end-over-end rotation for 4 h at 4°C. The beads were then washed four times with 1 ml IP-Lysis Buffer and bead-bound proteins were eluted by boiling in 1X Laemmli buffer (40 μl). Input and eluates were resolved by SDS-PAGE and subjected to Western blotting with a custom anti-Bre1 polyclonal antibody that was raised in rabbit or anti-Flag M2 (Sigma) antibody.

#### Protein stability assay

Yeast strains harboring *URA3*, 2μ plasmid with *GAL1* promoter-driven *Rad6-2Flag* or *rad6-A126T-2Flag* or *rad6-A126F-2Flag* were grown for 2 days at 30°C with constant agitation (230 rpm) in SC-URA with raffinose (2%) as the sugar source. After reinoculation and growth in fresh SC-URA with raffinose for 2-3 h, expression of Flag-tagged Rad6 or mutants was induced by adding 2% galactose and incubating with agitation for 2 h. Transcription was then shut-off by adding 2% glucose. Cells grown in raffinose or galactose media and at various time points after glucose addition were harvested for extract preparation using the TCA lysis method described previously(101). Briefly, log-phase yeast cells (20 × 10^7^) were harvested, washed once with phosphate-buffered saline (PBS) and once with 5% tricholoroacetic acid (TCA, Sigma) before storing at -80°C. Cell pellets were thawed in 20% TCA, lysed by bead beating and centrifuged (3000 rpm, 5 min at 4°C). The pellet was resuspended by vortexing in 1X Laemmli buffer (62.5 mM Tris.Cl pH 6.8, 10% glycerol, 2% SDS, 0.002% bromophenol blue, 2.5% β-mercaptoethanol). Subsequently the denatured lysate was neutralized by adding 2M Tris base before boiling for 8 min in a water bath and then clarified by centrifugation (13200 rpm, 10 min at 4°C). Protein concentration of the clarified lysate was measured using DC™ Protein Assay (Bio-Rad). Protein levels of wild-type or mutant Rad6 were determined by immunoblotting using anti-Flag M2 antibody (Sigma).

#### Immunoblotting

Yeast extracts were prepared using TCA lysis method as described above. Either equal amounts or a serial dilution of the lysates was prepared from various samples before resolving them in SDS-PAGE and transferring onto a polyvinylidene difluoride membrane. Following incubation with primary rabbit or mouse antibody and corresponding HRP-conjugated secondary antibody, protein signals were detected by chemiluminescence using Pierce™ ECL Plus Western Blotting Substrate (Thermo Scientific) and autoradiography. The following antibodies were used with their source and catalog numbers indicated within parentheses: anti-Flag M2 (F3165; Sigma), anti-Pgk1 (459250; Invitrogen), anti-GFP (AE011, Abclonal), anti-UBE2A (A7744; Abclonal), anti-UBE2B (A6315, Abclonal), anti-V5 (R690, Invitrogen), anti-H2B (39237; Active Motif), anti-H3 (ab1791; Abcam), anti-H3K4me1 (39297; Active Motif), anti-H3K4me2 (399141; Active Motif), anti-H3K4me3 (39159; Active Motif), anti-ubiquitin antibody (ab139467; Abcam); Mono- and polyubiquitinylated conjugates monoclonal antibody (FK2) (HRP conjugate) (BML-PW0150; Enzo Life Sciences), and anti-Rad6 (DZ33919; Boster Bio). Please note that the anti-UBE2B antibody recognizes both UBE2A and UBE2B (Supplementary Figure 2).

#### Nuclear magnetic resonance (NMR) spectroscopy

NMR data were collected on either a Varian INOVA 500 MHz using a room temperature HCN probe or a Varian INOVA 600 MHz equipped with an HCN Mark2 cryogenic probe. Data were processed and analyzed using NMRpipe(102) and Sparky(103) tools. Complete resonance assignment of wild-type Rad6 was accomplished using a standard suite of HCN triple resonance experiments (NHcoCA, HNCA, HNCACB, CBCAcoNH and ^15^N-edited NOESY) collected at two temperatures 25°C and 35°C with a uniformly labeled ^15^N, ^13^C, ^2^H (∼70%) Rad6 sample. Non-uniform sampling routines were used for all 3D HCN experiments(104). Isotope-labeled protein samples for wild-type or mutant yeast Rad6 or human UBE2B were prepared at 0.75-1.0 mM concentration in a buffer containing 25 mM sodium phosphate pH 7.5 and 0.5 mM TCEP. [^15^N, ^1^H] HSQC and HSQC-TROSY were recorded for yeast Rad6 and human UBE2B, respectively. Complete assignments for yeast Rad6 are at BMRB with accession code **<u>50964</u>**.

Chemical shift perturbations (CSPs) were qualitatively scored as follows: (1) wild type and mutant Rad6 or UBE2B were overlaid in Sparky. For each amide signal an overlap less than one-half the linewidth in either dimension was scored as CSP of 0, an overlap greater than one-half the linewidth and less than a full linewidth was scored as CSP 0.5, and an overlap greater than one linewidth was scored as CSP 1. CSP versus residue plots were generated using Graphpad Prism 9.0. CSP values were mapped on the crystal structure of yeast Rad6 (PDB ID: 1AYZ) (66) and human UBE2B (PDB ID: 2YB6) (63) and visualized using UCSF Chimera (105).

#### Molecular dynamics simulations

The crystal structures for yeast Rad6 (PDB ID: 1AYZ) (66) and its human homologs UBE2A (PDB ID: 6CYO) (80) and UBE2B (PDB ID: 2YB6) (63) were used to perform the classical molecular dynamics (MD) simulations using the GROMACS 2018.1 package (78). Amber99sb was selected as the forcefield for all the simulations (106). Models for the mutants A126T or A126F were prepared *in silico* using the crystal structures for Rad6, UBE2A or UBE2B in YASARA (107). The alanine was replaced with side chains from threonine or phenylalanine followed by a short minimization of 100ps. A freezing of all residues was performed except for those residues close to the point mutation to avoid any local crashes in the sidechains.

The nine systems (3 native and 6 mutants) prepared above were then solvated explicitly with TIP3P water molecules in a cubic box with a margin of 10 Å as previously described (108) and neutralized by adding sodium counter ions. Energy minimization using the steepest descent method for 5000 steps was carried out to remove any poor van der Waals’ contacts in the initial geometry. After the minimization step, two stages of equilibration were conducted: First, NVT (constant number, volume and temperature) equilibration was performed for 100 ps maintaining a constant temperature of 300K using V-rescale algorithm (109), with a coupling time of 0.1 ps and separate baths for the solute and the solvent. Second, NPT (constant number, pressure and temperature) equilibration was then performed with a constant pressure of 1 atm for 100ps using the Parrinello-Rahman pressure coupling scheme(110), with a time constant of 2ps. The position restrained NVT and NPT equilibration steps prompted water relaxation around the protein and reduced the system entropy. The covalent bonds were constrained by using the LINCS (Linear Constraint Solver) algorithm(111), and the electrostatic interactions were computed using the Particle Mesh Ewald (PME) method (112), with a cutoff distance of 10Å. A Lennard-Jones 6-12 potential was used to evaluate van der Waals interactions. Initial velocities were generated randomly using a Maxwell-Boltzmann distribution corresponding to 300 K. Finally, the production run was performed for 100ns for each prepared system without any restraints at 300 K in the isothermal-isobaric ensemble. A time-step of 0.002 ps was carried out in all the simulations and the MD trajectories were saved every 20 ps.

For trajectory analysis, the structural and conformational changes in the native and the mutant proteins were analyzed by applying *gmx rms* or *gmx rmsf* on trajectories resulting from the production run of simulations. Hydrogen bond interactions were quantified by *gmx hbond* tools of GROMACS program, and DSSP secondary structure evaluations and visualization were performed using VMD software (113). The minimized initial structure of each prepared system was used as reference geometry and all output files were analyzed and plots were created using XMGrace tool or Graphpad Prism 9.0. The simulations were repeated three times and the overall trajectories were similar between repetitions.

## Abbreviations used

ATP: adenosine triphosphate
PCNA: Proliferating cell nuclear antigen
MD: Molecular dynamics
RMSD: Root-mean-square deviation
RMSF: Root- mean-square fluctuation
DSSP: Definition of secondary structure of proteins
NMR: Nuclear magnetic resonance
HSQC: Heteronuclear single quantum coherence

## Data availability

Complete assignments for yeast Rad6 are at BMRB with accession code **<u>50964</u>**.

## Acknowledgements

We are grateful to Drs. Alexander Varshavsky, Rodney Rothstein, Brad Cairns, Tim Formosa, and the late Joe Horecka for kindly providing us plasmids or yeast strains. We thank Dr. Srividya Bhaskara for her valuable suggestions. We thank Kaitlin Radmall for technical assistance.

## Supplementary information

This article contains Supplementary information

## Tables

Table 1. Changes in folding free energy (*ΔΔG*) and predicted effect of mutations on protein stability obtained using Dynamut 2.0.

Table 2. Genotypes of yeast strains used in the study.

## Figure legends

**Supplementary Figure 1. Interactions with partner E3 ubiquitin-ligases are not impaired by Rad6 A126 mutations**. (a-b) *Left*, SDS-PAGE of lysates from uninduced (UIn) and IPTG-induced (In) bacterial cells, whole-cell lysates (WCL), and soluble and bound (B) fractions from cells coexpressing His_6_-Rad6 or His_6_-rad6-A126T or His_6_-rad6-A126F and *i*) Bre1R6BR or *ii*) GST-tagged Rad18R6BR. *R6BR*, Rad6 binding region. The *arrowhead*s indicate His_6_Rad6 or mutant. *Asterisks* indicate truncated product. *Right*, Histograms of average levels of copurifying Bre1R6BR or GST-Rad18R6BR normalized to the levels of partner wild-type or mutant Rad6 (two independent experiments). (c) Co-immunoprecipitation (IP) experiment. Immunoblots of lysates from yeast strains expressing Flag-tagged Rad6 or indicated mutants with 6HA-tagged Ubr2, 8V5-tagged Ubr1, or and 9Myc-tagged Rad18. The input was 5%of the lysate. Lysate from *bre1Δ* strain or strain expressing proteins without epitope tags served as controls for the anti-Bre1 and epitope-tag specific antibodies, respectively. The *asterisk* indicates cross-reacting protein. The arrow marked *LH* indicates immunoglobulin light chain.

**Supplementary Figure 2**. Time-course *in vitro* ubiquitin chain formation assay was performed using recombinant yeast Rad6 or human homologs UBE2A or UBE2B. Yeast or human Uba1 and ubiquitin were used with yeast Rad6 or its human homologs, respectively. Control reactions were performed without yeast Uba1 (*-E1*), without Rad6 (*-E2*), or without ATP. Blots were probed with antibodies recognizing ubiquitin, mono-or poly-ubiquitinated proteins (clone FK2), Rad6 or UBE2B. Note that the anti-UBE2B antibody recognizes UBE2A. *Ub*, ubiquitin; *Ub*_*2*_, diubiquitin; *Ub*_*n*_, ubiquitin chains or polyubiquitinated Rad6 or UBE2A or UBE2B; *E2-Ub*_*n*_, ubiquitinated Rad6 or its human homologs. Coomassie-stained gel shows the amount of Rad6, UBE2A or UBE2B used in each reaction.

**Supplementary Figure 3**. (a) Sequence alignment of Rad6 homologs, performed using Clustal Omega. Conserved residue (★), residues with strong (:) or weak similarity (.) are indicated. Uniprot IDs are indicated. *Arrows* indicate key residues of the catalytic pocket. The HPN motif and helix-3 are highlighted by *black lines*. (b) Superposition of structures of Rad6, UBC-1 and human Rad6 homologs. PDB IDs, key residues of the catalytic pocket and the position of A126 in helix-3 are indicated. (c) Alignment of the amino acids of helix-3 in yeast Rad6 and human homologs UBE2A and UBE2B. Position of the amino acids in the primary sequence are indicated.

**Supplementary Figure 4**. (a) RMSDs of Cα atoms of wild-type Rad6 (*red*), rad6-A126T (*black*), and rad6-A126F (*blue*). (b) Time evolution of secondary structural elements, based on DSSP classification, for wild-type Rad6 and its indicated mutants. *Arrows* indicate to secondary structures of mutants that were reorganized or disorganized at the end of the simulation in the mutant. (c) Analysis of the changes in the number of hydrogen bonds in Rad6 (*red*), rad6-A126T (*black*) and rad6-A126F (*blue*) over a 100-ns timescale. Three replicate simulations were run for these analyses.

**Supplementary Figure 5**. Histograms of the chemical shift perturbation (CSP) by residue for a) rad6-A126T and b) rad6-A126F relative to the structure of yeast Rad6. CSPs were scored for each mutant from the overlay of their NMR spectrum with that of wild-type Rad6 or UBE2B (see Figures 4b, 5b). No overlap of the NMR signal in a mutant relative to the wild-type was scored as 1, and partial overlap was scored as 0.5. The scores then were used to generate the histogram using Graphpad Prism 9.0. Schematic below the histogram shows the positions of various secondary structures

**Supplementary Figure 6**. (a) RMSD analyses of Cα atoms of UBE2A or UBE2B (*red*) or their mutants A126T (*black*), and A126F (*blue*). (b) Time evolution of secondary structural elements, per DSSP classification, for the indicated mutants of UBE2A or UBE2B. *Arrows* indicate secondary structures that were disorganized at the end of the simulation. (c) Analysis of the changes in the number of hydrogen bonds in UBE2A or UBE2B or their mutants over a 100 ns timescale. Three replicate simulations were run for these analyses.

**Supplementary Figure 7**. (a) Structures of UBE2A (PDB ID: 6CYO) and UBE2B (PDB ID: 2YB6) superposed onto the structure of UBC9 E2 from co-crystal structure of the UBC9-SUMO-RanGAP1 complex (PDB ID: 1Z5S) showing the location or proximity of helix-3 to catalytic pocket, ubiquitin and the incoming lysine of a substrate protein. (b) Zoomed image shows positions of key residues of the catalytic pocket of UBC9 and distances in angstroms (Å) from residues of the catalytic pocket to the glycyl-lysine isopeptide bond formed between SUMO and substrate RanGAP1.

