## SupplementalFigures for "Consequences of alanine-126 mutations in helix-3 on structure and functions of Rad6 E2 ubiquitin-conjugating enzymes"

### Supplementary Figure 1

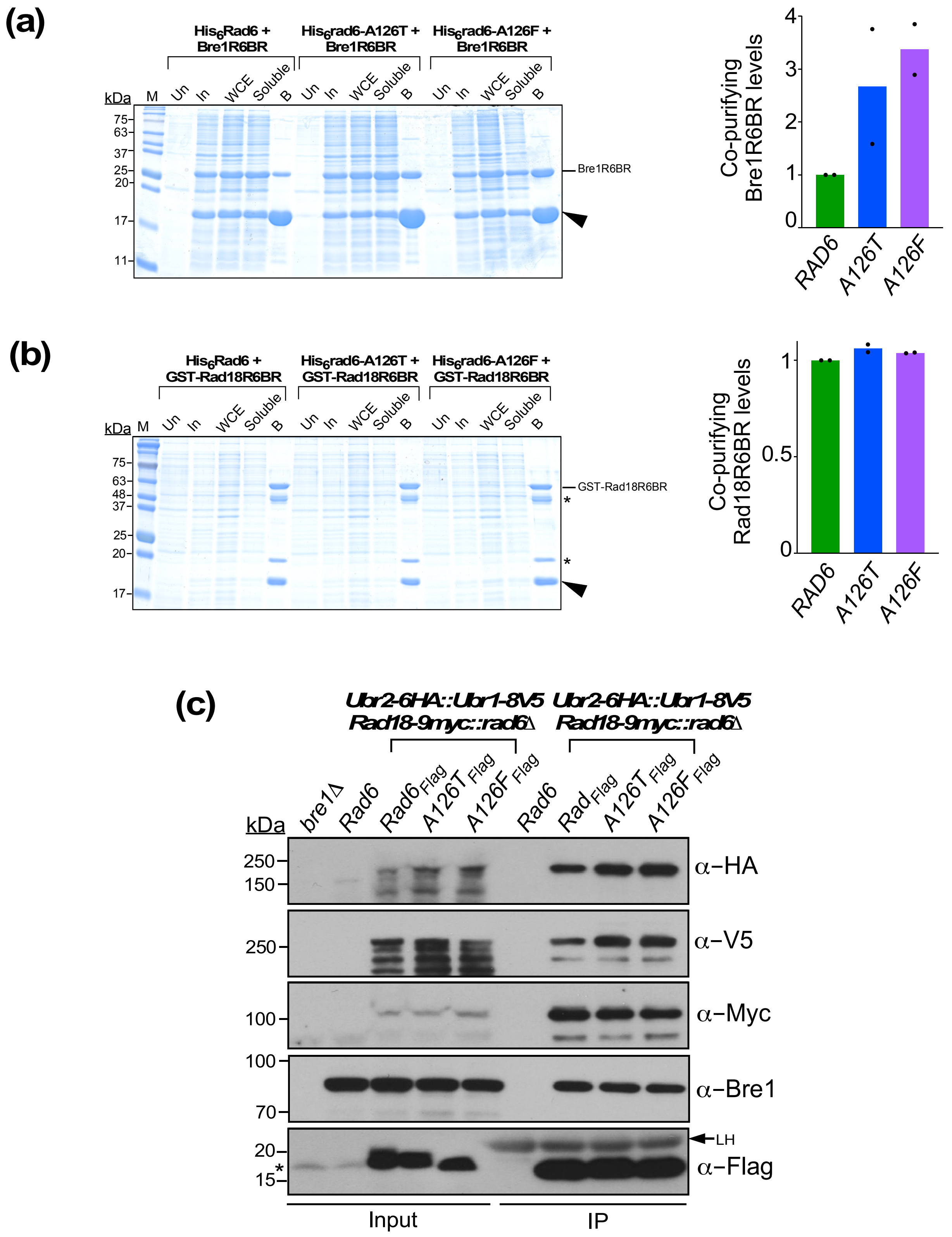

### Supplementary Figure 2

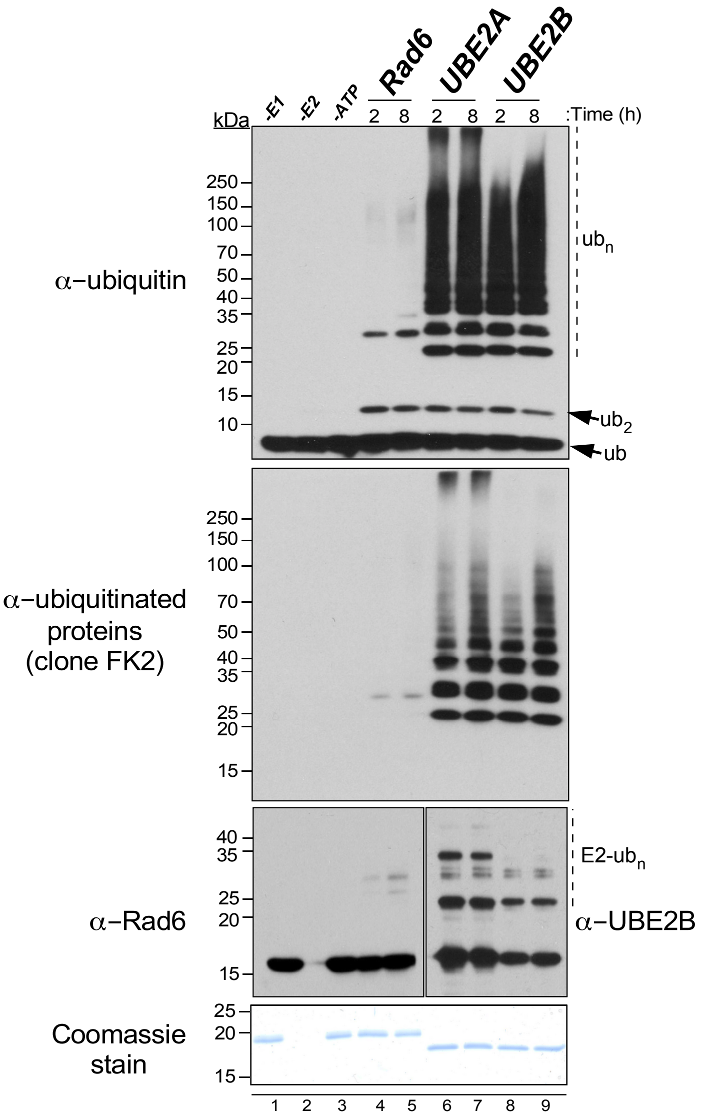

### Supplementary Figure 3

**(a)**

```
sp|P06104|UBC2_Scer     MSTPARRRLMRDFKRMKEDAPPGVSASPLPDNVMVWNAMIIGPADTPYEDGTFRLLLEFD
sp|P49459|UBE2A_Hsap    MSTPARRRLMRDFKRLQEDPPAGVSGAPSENNIMVWNAVIFGPEGTPFEDGTFKLTIEFT
sp|P63146|UBE2B_Hsap    MSTPARRRLMRDFKRLQEDPPVGVSGAPSENNIMQWNAVIFGPEGTPFEDGTFKLVIEFS
                        ***************::** * ***.:*  :*:* ***:*:** .**:*****:* :**

sp|P06104|UBC2_Scer     EEYPNKPPHVKFLSEMFHPNVYANGEICLDILQNRWTPTYDVASILTSIQSLFNDPNPAS
sp|P49459|UBE2A_Hsap    EEYPNKPPTVRFVSKMFHPNVYADGSICLDILQNRWSPTYDVSSILTSIQSLLDEPNPNS
sp|P63146|UBE2B_Hsap    EEYPNKPPTVRFLSKMFHPNVYADGSICLDILQNRWSPTYDVSSILTSIQSLLDEPNPNS
                        ******** *:*:*:********:*.**********:*****:*********:::*** *

sp|P06104|UBC2_Scer     PANVEAATLFKDHKSQYVKRVKETVEKSWEDDMDDMDDDDDDDDDDDDDEAD
sp|P49459|UBE2A_Hsap    PANSQAAQLYQENKREYEKRVSAIVEQSWRDC--------------------
sp|P63146|UBE2B_Hsap    PANSQAAQLYQENKREYEKRVSAIVEQSWNDS--------------------
                        *** :** *::::* :* ***.  **:**.*
                            α−helix-3
```

**(b)**

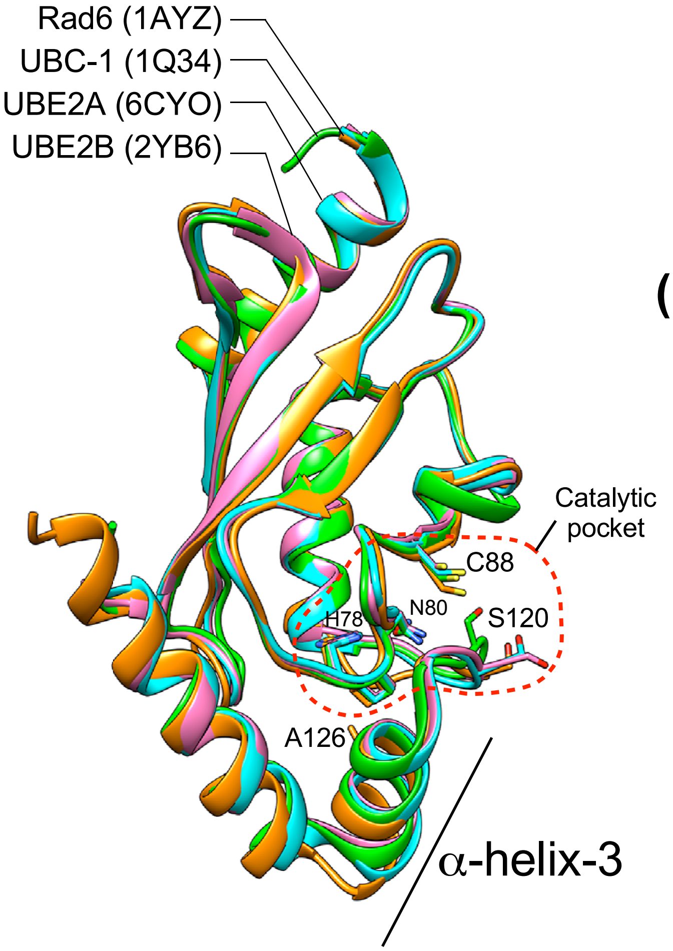


**(c)**

```
            124 125 126 127 128 129 130 131 132
Rad6:  ..VEAATLFKD..
UBE2A: ..SQAAQLYQE..
UBE2B: ..SQAAQLYQE..
            :**  *:::
           α−helix-3
```

### Supplementary Figure 4

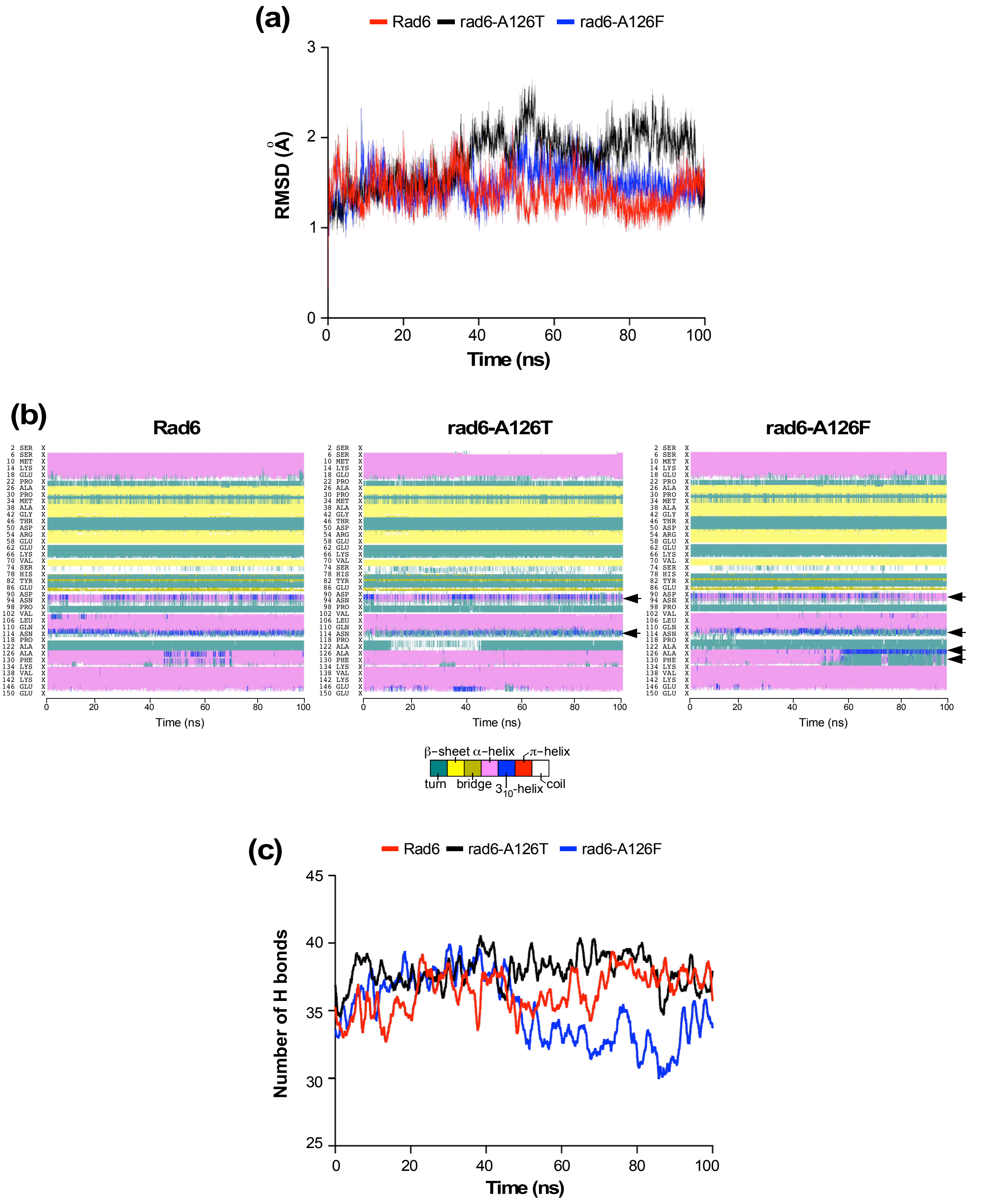

### Supplementary Figure 5

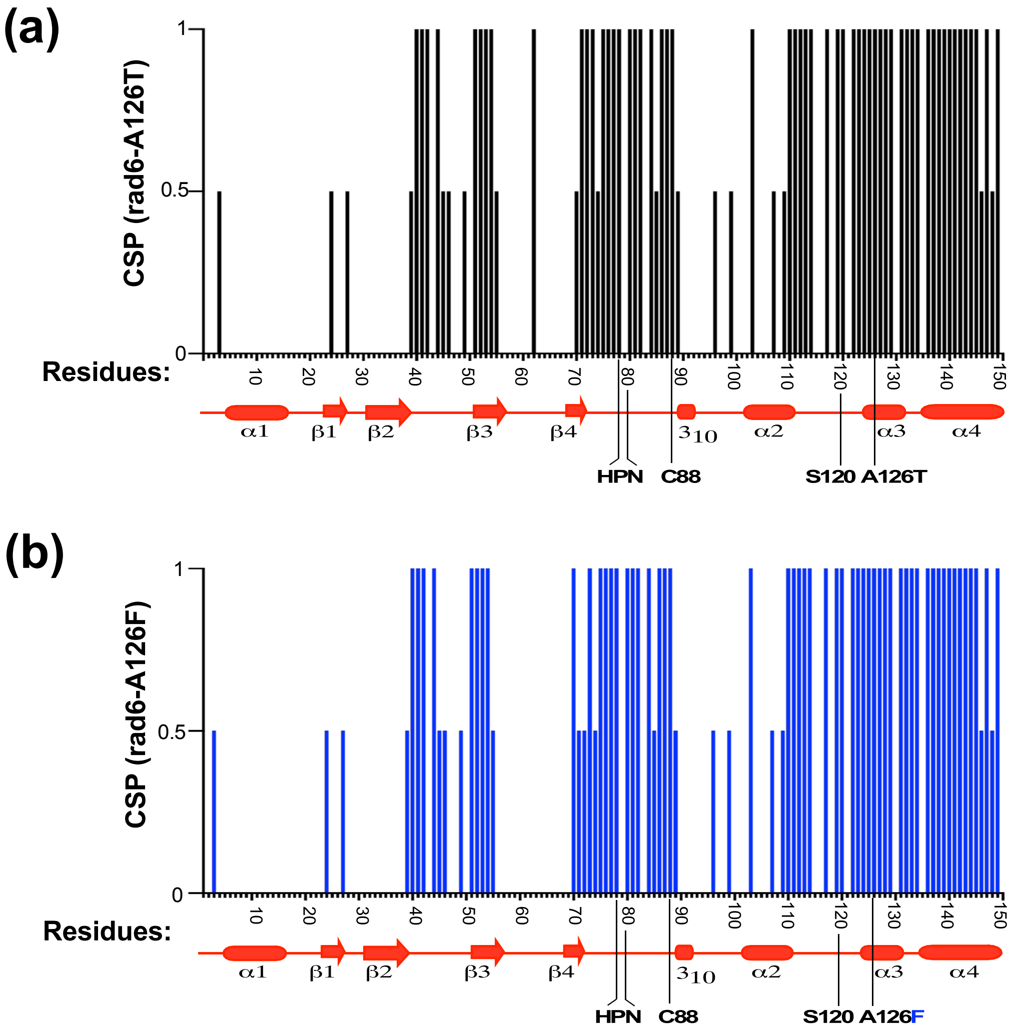

### Supplementary Figure 6

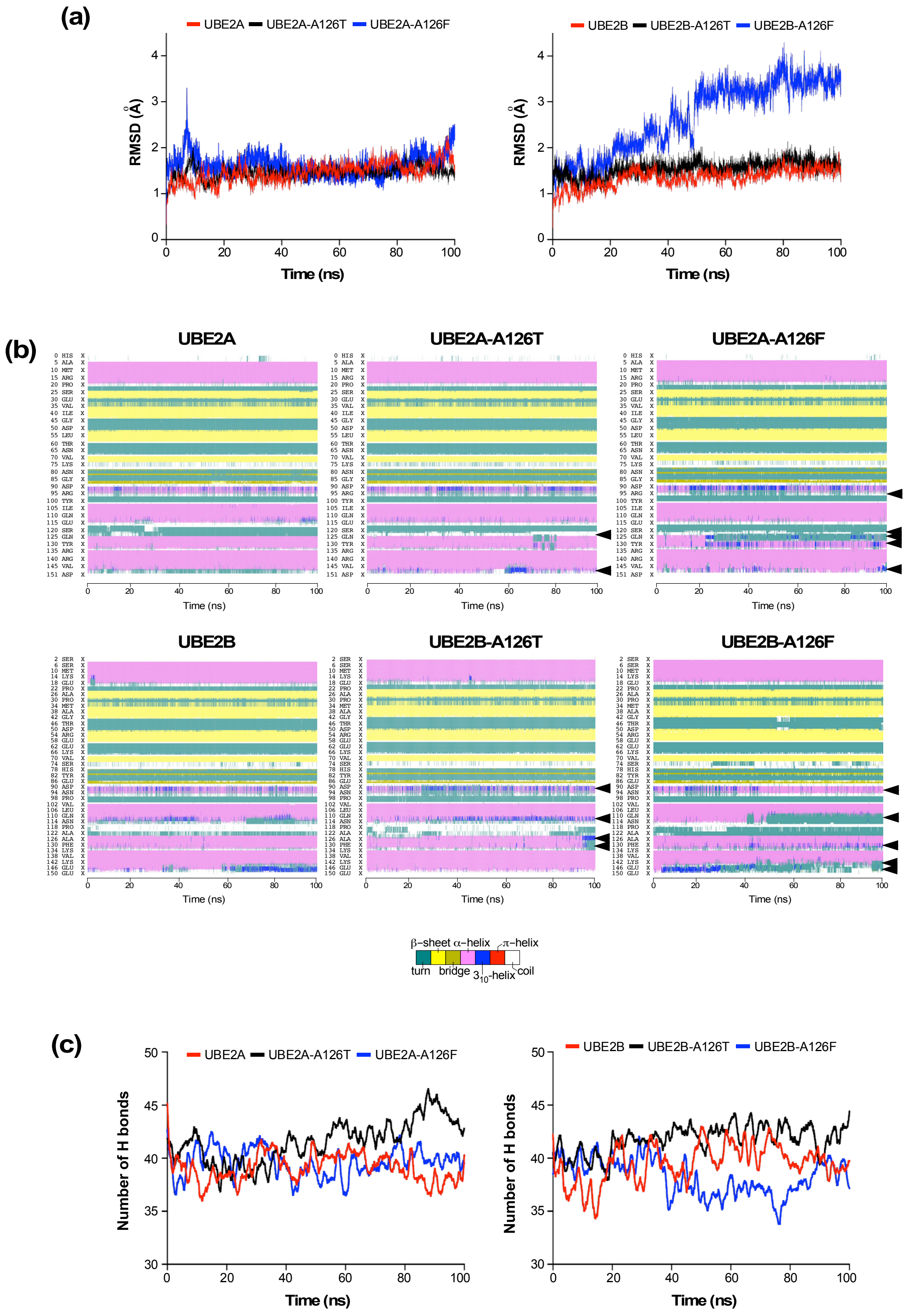

### Supplementary Figure 7

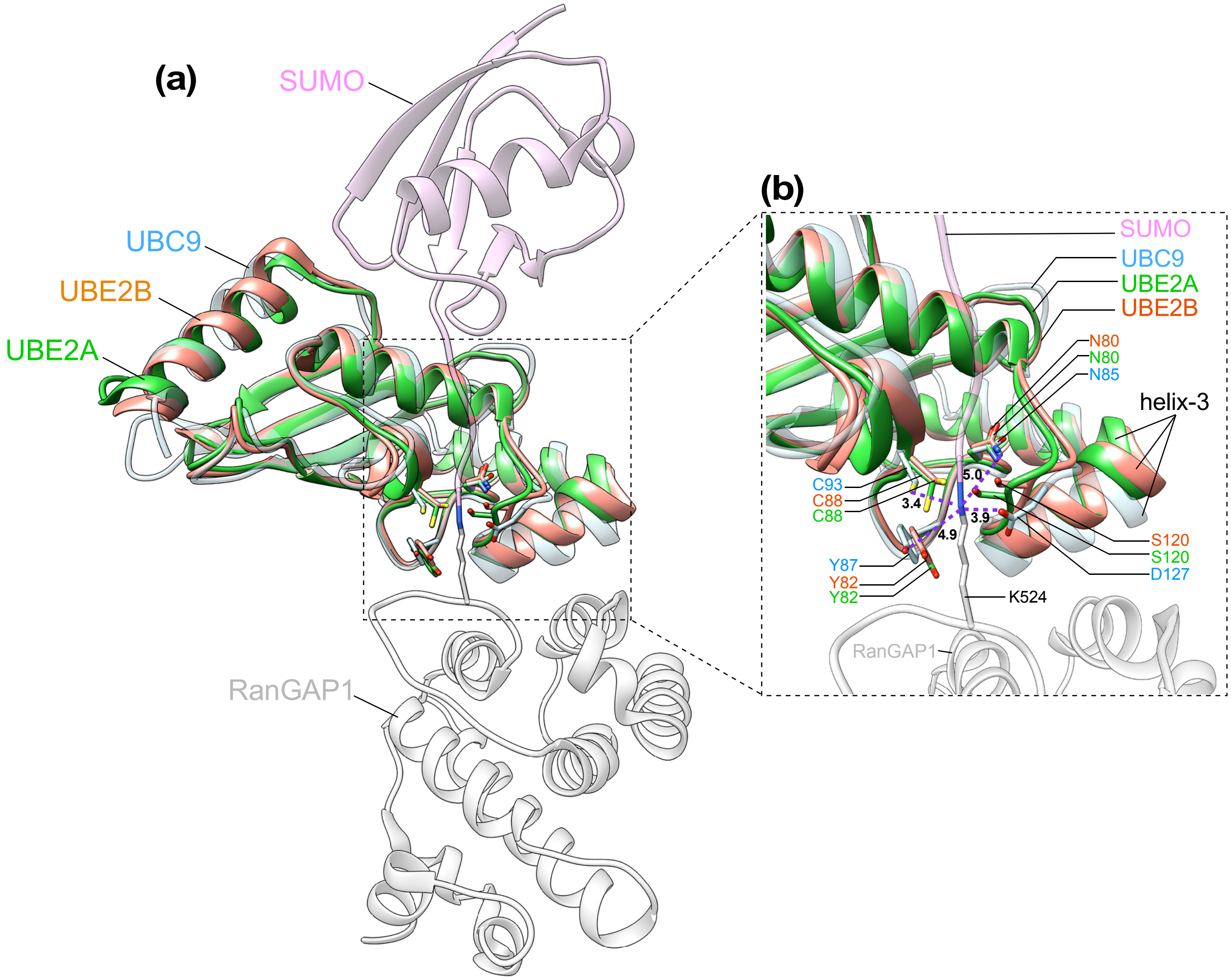
